# It’s All In The Journey: Putative Strategies Extracted from Navigation Paths Predict Spatial Memory and Hippocampal Recruitment

**DOI:** 10.64898/2025.12.14.693950

**Authors:** E. Zita Patai, David Stawarczyk, Nora Herweg, Carlos A. Gomes, Hui Zhang, Andreas Schulze-Bonhage, Lukas Kunz, Nikolai S. Axmacher

**Affiliations:** Department for Neuropsychology,Institute of Cognitive Neuroscience, Ruhr University Bochum, Germany; Psychology and Neuroscience of Cognition Research Unit, University of Liège, Place des Orateurs 1 (B33), 4000 Liège, Belgium; GIGA – CRC (Cyclotron Research Center) Human Imaging, University of Liège, Allée du 6 Août 8 (B30), 4000 Liège, Belgium; Epilepsy Center, Faculty of Medicine, University of Freiburg, Germany; Department of Epileptology at the University Hospital Bonn, Germany

## Abstract

Navigation in the real world may rely on a multitude of different strategies with distinct advantages and disadvantages. However, the characterization of spontaneous strategy use during wayfinding and underlying neural correlates are scarce. We explore data across three groups of participants who performed a virtual navigation task: (I) epilepsy patients undergoing invasive electrophysiological recordings of the hippocampus, and two groups who underwent fMRI scanning: (II) young adults at genetic risk for Alzheimer’s disease (APOE-e4 carriers) and (III) healthy controls. We establish three metrics by which to quantitatively characterize each taken path, with the aim of simultaneously reflecting the putative strategies used by humans to solve the task: 1. straightness of paths; 2. deviation towards the edge of the circular arena; and 3. overlap of individual paths leading to a specific object location, which putatively relate to reliance on allocentric (1) and egocentric reference frames (2 & 3). Extracted metrics were related to performance in the task and to hippocampal activity. We find that strategy use is variable within participant, changes across the experiment, and is adaptive in a group-dependent manner. While hippocampal BOLD activity and theta (gamma) power were generally inversely (positively) related, we found that the recruitment of the hippocampus was both group- and strategy- specific. Our results further our understanding of the heterogeneous and idiosyncratic nature of spatial navigation by revealing, how these behaviors are altered by disease and the subsequent compensatory neural dynamics, as well as the link between mesoscopic and macroscopic neural signals.

## Introduction

Behavior is variable and complex, and often not what the experimenter expects – even to solve a simple task of getting from A to B, humans can employ different strategies to get to their goal [Ekstrom et al., 2018, Hegarty et al., 2023]. Despite being in the same environment, we may either choose to use local landmarks and boundaries to guide our path, memorize a series of turns, or use distal landmarks to form a metric representation of our surroundings – a so-called cognitive map [Tolman, 1948]. Since the discovery of place cells [O’Keefe and Dostrovsky, 1971] and the seminal paper demonstrating the hippocampal role in place as contrasted with response learning [Packard and McGaugh, 1996], the hippocampus has been proposed to be the neural locus of the cognitive map [Epstein et al., 2017, McNaughton et al., 2006, Behrens et al., 2018, Moser et al., 2008]. Other neural signals in the medial temporal lobe, in the form of grid cells, border cells, and head direction cells etc. contribute to the ability to successfully navigate space by storing spatial representations in distinct formats [Grieves and Jeffery, 2017]. This understanding of different neural signals in the brain, in turn, has contributed mechanistic evidence on the differentiation of self-referenced (egocentric) vs. world-referenced (allocentric) representations or reference frames [Ekstrom et al., 2014, Buzśaki and Moser, 2013],but see [Filimon, 2015]. The former relies on self-referenced motion signals or orientation to local landmarks and boundaries, and the latter on understanding of the relationship between environmental cues (e.g. distal landmarks). Allocentric representations are independent of the observer’s position and purportedly reliant on the hippocampus and related medial temporal lobe structures, forming what is known conceptually as the cognitive map.

Many paradigms have been developed to tease apart the contribution of these neural representations, and the putative reference frames they support, by controlling experimental variables to force the participant into using either an allocentric or egocentric strategy [Igĺoi et al., 2009, Grech et al., 2018, Nett et al., 2025, Marchette et al., 2011]. However, real-world navigation behavior may combine multiple strategies, and neural representations may not be exclusive. It thus appears surprising that although humans and animals show heterogeneous navigation strategies at individual [Hegarty et al., 2023] and cultural levels [Fernandez-Velasco and Spiers, 2024], little research has examined how strategy use develops spontaneously in open or unrestricted environments and how this can be quantified. This differs from other fields, such as those investigating migratory birds, which have adopted trajectory analysis techniques to describe large-scale movement patterns in space (e.g., [Tao et al., 2021]). One notable exception is the approach by [Garthe et al., 2009], in which the search paths taken by mice in a Morris water maze were categorized based on various factors, including direction and amount travelled, parts of the arena explored, and perseverance. The authors found that search strategies changed over the course of exposure to the maze, became more indicative of flexible (i.e., allocentric) spatial representations, and that suppression of adult neurogenesis in the hippocampus altered this pattern. A more recent study quantified a metric (“target estimation vector”) that captured the animal’s route in terms of distance and direction from a goal location, and showed that direction information was learned earlier than distance information, but no neural effects were reported [Xu et al., 2024].

While not directly characterizing the movement paths themselves, others have found evidence that ongoing hippocampal activity relates to behavioral and cognitive aspects of navigation (for parsimony, we focus on human experiments largely inspired by animal work). Foremost, this relates to the ubiquitous theta rhythm in the hippocampus, oscillating around 4-8Hz [Buzśaki, 2002, Bohbot et al., 2017]; in humans also lower and broader frequencies have been reported, [Jacobs, 2014, Guth et al., 2025]. Theta oscillations have been associated with travelling speed [Aghajan et al., 2017, Goyal et al., 2020, Watrous et al., 2011], distance travelled and distance to goal location [Bush et al., 2017, Liu et al., 2023, Vass et al., 2016], distance to boundary [Maoz et al., 2023, Stangl et al., 2021] and the differentiation of individual goal representations [Kunz et al., 2019b, Wang et al., 2018] during spatial navigation using invasive recordings. Functional magnetic resonance imaging (fMRI) in healthy humans has shown that the hippocampus codes for ongoing distance to navigation goals [Bierbrauer et al., 2020, Patai et al., 2019] for review see [Spiers and Barry, 2015, Nyberg et al., 2022], allocentric spatial and temporal distances [Deuker et al., 2016], and spatial topography of the environment [Javadi et al., 2017].

These studies provide insight into hippocampal activity during spatial processing, both for egocentric and allocentric coding schemes. However, the variables above have often been explored in isolation and are somewhat removed from the more autonomous or decision-related aspects of navigation – specifically, how humans choose to solve navigation tasks when unconstrained in the paths they can take. They do highlight that hippocampal coding is dimensional rather than categorical, as both theta power and blood-oxygen level dependent (BOLD) activity scale with the magnitude of these variables. Furthermore, evidence suggests that neural signals may not be tuned to only one aspect of the environment but can support multiple functions, rather than being just ‘place’ or ‘grid’ cells [Burgess et al., 2005, Hardcastle et al., 2017b, Hardcastle et al., 2017a, Stefanini et al., 2020]. Thus, it stands to reason that this high representational dimensionality may also be reflected in behavior. A more direct approach would be to quantify, across multiple dimensions (or putative strategies), the characteristics of paths taken in an environment, to understand how these relate to ongoing hippocampal activity (rather than relying on explicit categorization). This is what we aimed for here, considering possible changes in linear relationships between strategies, performance, and neural activity by adopting piecewise regression analysis [Muggeo, 2003].

Moreover, only few human studies (e.g. [Kaplan et al., 2012]) have examined the relationship between markers of navigation across different recording methodologies (invasive/non-invasive electrophysiology and functional imaging). Further, navigation-related behaviors often vary across paradigms (including experimental constraints, designs, and human idiosyncracies), thus making comparisons difficult between studies. Even within one methodological approach (e.g., fMRI), subtle changes in the paradigm may completely invert the relationship between hippocampal signals and, for example, representation of distance [Spiers and Barry, 2015]. We still have limited understanding of how oscillations measured via local-field potentials (LFPs; such as during intracranial recordings of epilepsy patients), and macroscopic brain activity, measured via functional imaging (BOLD) signals, are related. Although it is generally assumed that high frequency oscillations are positively, and low frequency oscillations negatively, related to BOLD activity (for review see [Kunz et al., 2019a]), only one study has directly compared theta power and parahippocampal BOLD activity, finding a positive relationship [Ekstrom et al., 2008]. To the best of our knowledge, no previous study examined within the same paradigm how hippocampal theta oscillations, the ‘hallmark of navigation’, are related to changes in hippocampal BOLD activity, which are not only indirect, but measured on a time scale an order of magnitude longer [Ekstrom, 2010].

How might one go about characterizing paths in a quantitative way? Cognitive map-based navigation can be characterized by the ability to perform short-cuts (as demonstrated in bats:[Harten et al., 2020, Toledo et al., 2020]; mice: [Xu et al., 2024]; humans: [Marchette et al., 2011]). Short-cuts can be defined as the shortest paths from a variable start location to a goal (a straight line in a circular arena), rather than repeating previously taken routes. Accordingly, one way to quantify cognitive map-based navigation is by measuring how straight or direct a path is (allocentrically). On the other hand, egocentric approaches would rely on orienting relative to landmarks and taking previously experienced routes. This can be quantified by the percentage of overlap in the x-y coordinates of paths (stereotypicality or route matching).

Individuals carrying the APOE-e4 allele, the most important genetic risk factor of late-onset Alzheimer’s disease [Corder et al., 1993], have been shown to preferentially approach environmental boundaries and landmarks during spatial navigation tasks [Bierbrauer et al., 2020, Coughlan et al., 2019, Kunz et al., 2015]. This behavior may be conceptualized as an attempt to integrate egocentric and allocentric reference frames, as boundaries can serve as local landmarks, primarily providing distance rather than directional information (anchoring). In fact, in early reports using the virtual arena task [Doeller and Burgess, 2008], selective removal of the environmental boundary caused impaired performance despite the presence of distal landmarks, which consistently provide directional information—further emphasizing that boundaries support learning locations relative to surface geometry. Rats in the Morris water maze also show a tendency to move towards the boundaries as well as to use distal cues (configural information) to solve the task [Maurer and Derivaz, 2000].

However, apparent employment of each of these strategies may also reflect poor performance; for example, participants may choose to move close to the boundary when they lost orientation, or move on straight lines when uncertain about the goal location, and these movements might then be associated with relatively diffuse and imprecise internal representations of space. Thus, the relationship of movement patterns to performance might be nonlinear.

Given the wealth of data obtained during navigational tasks, both at the behavioral and neural level, we chose to reanalyze three existing datasets previously reported in the literature [Chen et al., 2018, Liu et al., 2023, Kunz et al., 2015, Kunz et al., 2019b]. We explore data across three groups of participants: I) epilepsy patients undergoing invasive recordings of the hippocampus, and two groups who underwent fMRI scanning: II) young adults at genetic risk for Alzheimer’s disease (APOE-e4 carriers) and III) healthy controls. All participants performed an identical spatial navigation task, and we characterized each of the paths that participants spontaneously took, trial-by-trial, across three metrics: how straight they moved, the amount they deviated towards the boundary, and how much they used overlapping paths (Figure 2A). Taken together, these metrics provide insight into the multidimensionality of chosen paths and strategies when navigating to hidden goal locations.

We aim to address three outstanding questions in the literature: Firstly, how the use of different, non-exclusive, spontaneous strategies to arrive at a goal location relates to performance as well as to ongoing neural oscillations (measured via intracranial EEG in patients with epilepsy) and to BOLD activity (measured via fMRI in both APOE-e4 carriers and healthy young adults) in the hippocampus as well as two control areas, the entorhinal cortex and the caudate nucleus. Second, as APOE-e4 carriers show no overt performance deficits in this task [Kunz et al., 2015], we investigate if this is related to the employment of alternative strategies and potential compensatory neural mechanisms [Cabeza et al., 2018, Trachtenberg et al., 2012]. Third, given the identical nature of the task across different methodologies, we aim to understand how neural signals in the hippocampus represented by mesoscopic (iEEG theta oscillations) and macroscopic (BOLD) measures are related to each other during ongoing behavior.

## Results

We analyzed data from 36 Controls, 38 APOE-4 carriers (“APOE-e4”), and 33 epilepsy patients with intracranial hippocampal EEG electrodes (iEEG) while they performed a navigation task in which they had to find multiple hidden object locations in a circular arena (Figure 1A). We defined performance as Drop Error, i.e. the distance between the indicated and the actual object location in a given trial. To correct for the fact that Drop Error could by chance be larger for objects located closer to the boundaries than those at the center of the arena, we estimated a “corrected” Drop Error metric, which was defined as the percentile rank score of Drop Error on every trial, compared to a distribution of random drops around the arena (a larger number indicates worse performance; See Methods, [Kunz et al., 2021, Kunz et al., 2024, Miller et al., 2018]. A Kruskal-Wallis test showed Drop Error was significantly different across groups (H(2,107)= 53.123, p *<* 0.001), and Wilcoxon rank sum tests showed Drop Error was higher in the iEEG group than both the Control (W=70, p*<* 0.001,r=0.76) and APOE-e4 group (W=71, p*<* 0.001,r=0.76), while the APOE-e4 carriers and Controls did not differ in their performance (p=0.76, Figure 1B).

**Figure 1:**
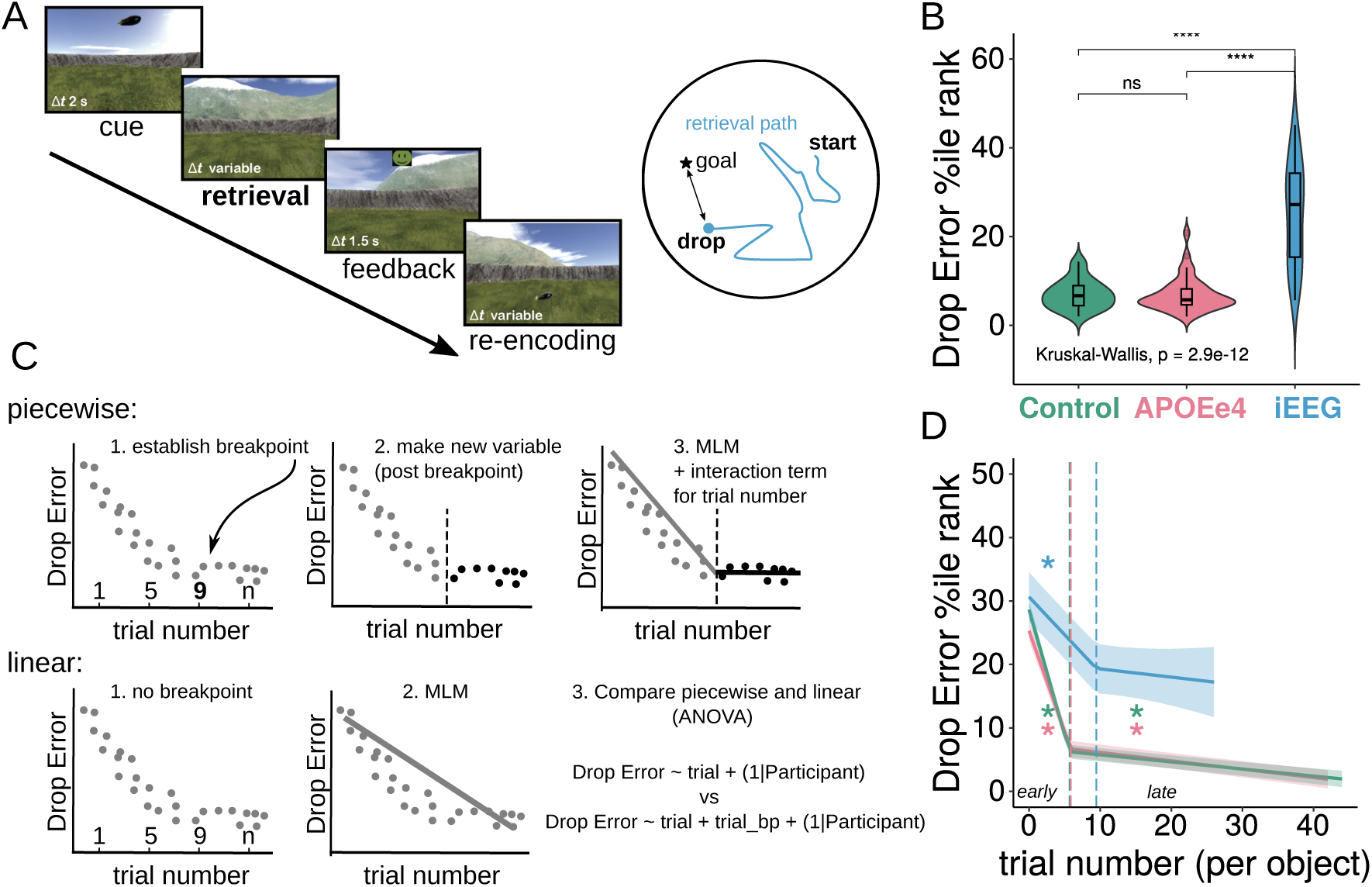
Navigation Task and Performance. A) Left: Schematic of a trial in the task (note: for the iEEG patients, the cue period only showed the object for 2s, not the environment (black background)). Right: Schematic of arena overview with a hypothetical start location and path leading to the drop location, as well as the goal location. Participants took part in a desktop virtual navigation task and searched for hidden object locations within a circular arena. After an object was presented (“Cue”), participants navigated through the arena (“Retrieval”) and indicated when they believed to have reached it (drop location). They were given feedback as to their performance and then were shown the correct object location (goal location) to which they navigated to start the next trial (“Reencoding”). Right: During retrieval, we recorded participants’ paths. Start location was the goal location from the previous trial (the first trial started from a random location). Performance was measured via the distance between the goal location and the drop location (“Drop Error”). B) Drop Error (percentile rank against chance, i.e. corrected for the bias in potential error as a function of object location, see Methods) differed significantly between groups, with epilepsy patients (iEEG) performing significantly worse (higher Drop Error) than both Control and APOE-e4 groups (both p*<* 0.001). C) Schematic of piecewise regression, used throughout the study and compared to linear regression. D) Drop Error decreased as a function of trial number per object. Empirically defined breakpoints revealed that a rapid decrease of Drop Error in the early trials was followed by a slower decrease afterwards (‘early’ and ‘late’ periods). Vertical lines indicate significant breakpoints. Stars above the regression lines indicate whether they are significant, after FDR correction. In the case of the iEEG group, both early learning slope and the breakpoint are significant, but the second leg of the regression (i.e. the slope in late learning) is not, indicating that after about 9 trials of an object, performance plateaued. Controls and APOE-e4 carriers showed steep learning early and continued (though significantly slower) learning after the breakpoint.

Given that participants were presented with the correct object location at the end of each trial, we hypothesized that Drop Error would decrease as a function of learning, i.e. as increasing trials per object location were experienced. We observed that learning seemed to be non-linear and opted for piecewise linear models as they are easier to interpret cognitively and less prone to over-fitting than polynomial and/or generalized additive models, and ran these effects models for each group (see Figure 1C and Methods for details) to investigate Drop Error as a function of trial number. Briefly, these models first establish a breakpoint in the slope between the outcome variable (in the example, “Drop Error”) and the predictor variable (“trial number”). A new variable is then created, to capture the data after this breakpoint. This variable is zero before the breakpoint, and coded as “value minus breakpoint” thereafter (e.g., if the breakpoint is trial 9, the value at trial 10 is 1; see Figure 1C). A mixed-linear model was run with this new variable included as a fixed effect (as well as the original predictor variable, in this case, trial number). In all subsequent analyses, the piecewise linear model was statistically compared to a linear model (tested via ANOVA, see Methods for details) and reported only if it outperformed the latter. In the reported results, the first beta estimate in these models relates to the slope and significance before the breakpoint and the second indicates whether the post-breakpoint slope is different from the first. A separate test is used to test whether this second slope (which is a sum of the two beta values) is significant itself (see Methods for details).

We found in all groups a steep decrease of Drop Error during the early periods of the experiment, indicating fast learning (Figure 1D, 7 trials per object in the Control and APOE-e4 group, and 9 trials per object in the iEEG group, Supplemental Table 7 for details), followed by a significantly less steep decrease (i.e. slower learning) in the remaining trials. For subsequent analyses, we thus divided navigation behavior into ‘early’ and ‘late’ learning phases.

### Definition of putative strategies via path metrics

We quantified navigation strategies via three metrics that compared in each trial, the path taken by the participant (“observed”) to the possible “direct” paths between the start location and the drop location indicated (Figure 2A). The direct paths between the two points are considered in order to normalize the metrics to a ground truth, i.e., to account for the possibilities in movement given the locations of start and drop locations within the arena, and thus for any goal location-specific differences that could contribute to changes in strategy across trials and/or participants.

**Figure 2:**
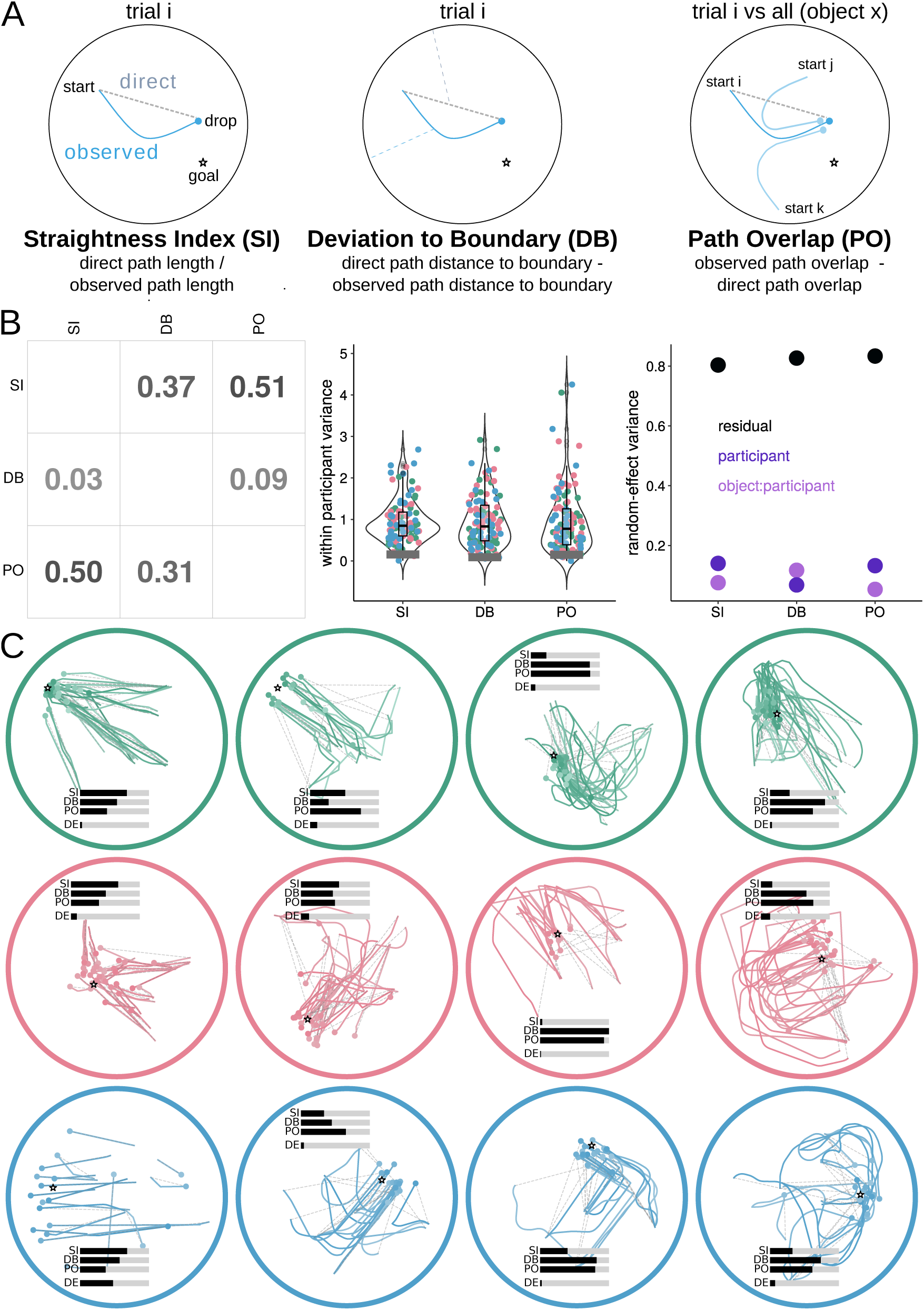
Quantification of movement paths via putative strategies. A) Schematic of three path metrics computed for each trial. Straightness Index (SI) was taken as the ratio between the length of the direct path and the length of the actual path [allocentric]. Deviation to Boundary (DB): Difference between the distance of the direct and observed paths from the boundary, respectively. A positive value indicates that the observed path deviates towards the boundary, a negative that the observed path deviates towards the center [anchoring]. Path Overlap (PO) is the difference between the observed path overlap (of a given trial to all other trials for that particular goal object) and direct path overlap (of a given trial to all other trials for that particular goal object) gives a measure of how stereotypical each path was when approaching a specific goal location [route matching]. B) Left: marginal r2 values from piecewise regressions where each metric was used to predict each other metric (outcome variable on y axis, predictor on the x axis). Metrics were related but explained no more than half of the variance between them, and were not linearly related, see Supplemental Figure 1. Middle: High variability of strategy use across trials as can be seen from the within-subject variance plotted for three metrics (color coded by group), with between-participant variance indicated by the grey bar. Right: linear mixed-effects model (simple intercept model only) showing the same pattern, where residual (i.e. within-participant) variance is much larger than both random effects of participant as well as objects (which are nested within participants). C) Path examples with metrics. Each panel shows the observed paths (colored lines) taken by a participant towards a particular goal location (indicated by a star), as well as the direct paths (grey dashed lines). Inset are the z-scored metrics, shown on a slider scale for easier comparison across metrics (scale: z=–2 to 2), along with mean corrected Drop Error (scale: DE = 1-50) for the goal location shown. For this display, we have also averaged the trial-level measures of Straightness, Deviation to Boundary and Path Overlap metrics across the goal location presented. The color of the circle indicates group membership (green = Controls, pink = APOE-e4, blue = iEEG). From left to right: examples of high straightness trials, paths that are deviating to the center or the boundary, and varying levels of path overlap on the goal approach.

First, in order to quantify the existence of a “cognitive map” of the environment and relationship between goal locations (putatively an allocentric strategy), we extracted the Straightness Index (SI). This was defined as the ratio between the length of the direct path and the length of the actual path. Values closer to 1 would indicate straighter paths. Second, we analyzed how much participants used the boundary for orientation, we extracted how much they deviated towards the boundary (putatively an “anchoring” strategy, which could indicate reliance on a combination of allocentric and egocentric strategies). This was quantified as the difference between the distance of the direct paths from the boundary vs. the distance of the observed paths from the boundary. A positive value indicates that the observed path deviates towards the boundary, a negative that the observed path deviates towards the center. This metric was measured trial-by-trial by taking the median of the deviation values across the whole path (DB). Third, we investigated the degree to which participants approached drop locations via stereotypical paths with matched viewpoint (putatively an egocentric strategy). Path Overlap (PO) was calculated as the percentage of overlap in x/y coordinates of a path in a given trial with all other trials taken to a particular goal location.

This measure was again normalized by the overlap of the direct path to all other trials. A positive value would indicate that the observed path to the drop location overlapped more with the paths from the other trials involving the same goal location than the direct path did. All metrics were z-scored to facilitate comparison across metrics (calculated across all trials from all participants, see Methods for details).

These three metrics were significantly though non-linearly related – for example, straighter paths were associated with reduced deviation to either the boundary or the center (Supplementary Figure 1 and Supplementary Table 5). Importantly, however, employment of these strategies was neither mutually exclusive nor did any of the metrics explain the variance in any other metric by more than 51%, as quantified in piecewise regressions where metrics were used as outcome or predictor variables (Figure 2B left panel, also see Supplemental Figure 5 for details). These relationships highlight that navigation trajectories can be characterized across multiple metrics simultaneously (see also Figure C for examples), and that they do not necessarily have to be categorized as belonging to one strategy over another (e.g.: egocentric vs allocentric).

We next tested whether strategy use reflects a stable trait measure or whether it differs substantially across trials. We therefore analyzed the variance in strategy use across trials in individual participants and compared this to the variance in average strategy use between participants. We found that the average within-subject variance of strategy use was much higher than the between-subjects variance of subject-averaged strategy use (all within-subject variance *>* 0.9, compared to across-subject variance all *<* 0.6, Wilcoxon signed-rank tests all p*<* 0.001, Figure 2B middle panel). We confirmed this via measuring the random effects variance in mixed effects model (simple intercept model only) and showed that residuals accounted more than fourfold the variance as compared to random effects such as participant or object identity (Figure 2B right panel; ICC values were between 0.18-0.21 for all strategy metrics, further underscoring that the majority of variability comes from within the grouping variables, i.e. within participant). These findings strongly suggest that strategies are spontaneously adopted to a variable degree in individual trials, possibly depending on a multitude of factors, rather than being a characteristic at the individual participant level.

Finally, we found that these metrics were significantly different when comparing paths from retrieval and re-encoding phases, i.e. when the goal object was shown to participants (increased SI and reduced DB/PO in re-encoding, see Supplemental Figure 2), emphasizing that these are strategies employed selectively during memory-based spatial navigation, rather than during movement per se (or ‘guided navigation’) within the arena.

### Group differences in strategy use

To understand how the three groups may differ in their use of these putative strategies, we averaged the path metrics across all trials within participants (Figure 3A). SI did not differ significantly between groups (Kruskal-Wallis test: H(2,107)=2.79, p=0.25). DB differed between groups (H(2,107)=18.16, p*≤* 0.001). Compared to Controls, both APOE-e4 carriers (Wilcoxon rank sum tests: W=957, p=0.009, r=0.33) and iEEG patients (W=249, p*≤* 0.001, r=0.5) deviated more to the boundary, and iEEG patients deviated by trend more to the boundary than APOE-e4 carriers (W=522, p=0.069, r=0.14). When compared to a baseline of zero (no deviation), Controls actually deviated significantly towards the center of the arena (W=185, p=0.02), while iEEG patients deviated significantly towards the boundary (W=507, p*≤* 0.001) and the APOE-e4 group showed a trend (W=489, p=0.09). Relatedly, APOE-e4 and iEEG groups showed a significant bias towards indicating drop locations closer to the boundary (both p*≤* 0.001), whereas Controls showed no such bias in drop location (‘Drop Error boundary bias’, also a significant main effect of group (H(2,107)=15.25, p*≤* 0.001, Supplemental Table 1). These results corroborate previous findings that APOE-e4 carriers tend to navigate closer to boundaries/landmarks [Kunz et al., 2015, Bierbrauer et al., 2020] and show similar effects in iEEG patients who often have dysfunction of MTL structures (see Discussion). PO differed between groups as well (H(2,107)=12.74, p=0.002), with Controls (W=875, p=0.002, r=0.4) and APOE-e4 carriers (W=817, p=0.08, r=0.26) using more overlapping paths than the iEEG group. These findings suggest that APOE-e4 carriers and in particular iEEG patients employ more “anchoring” strategies than Controls, and the reverse pattern was found for egocentric “route matching” strategies. Putative strategies did not differ depending on gender and no age effects were found (except in the APOE-e4 group, increased age was related to increasingly straight paths and less route matching, but note the age range was only 18-29 years of age in this group; see Supplemental Table 4).

**Figure 3:**
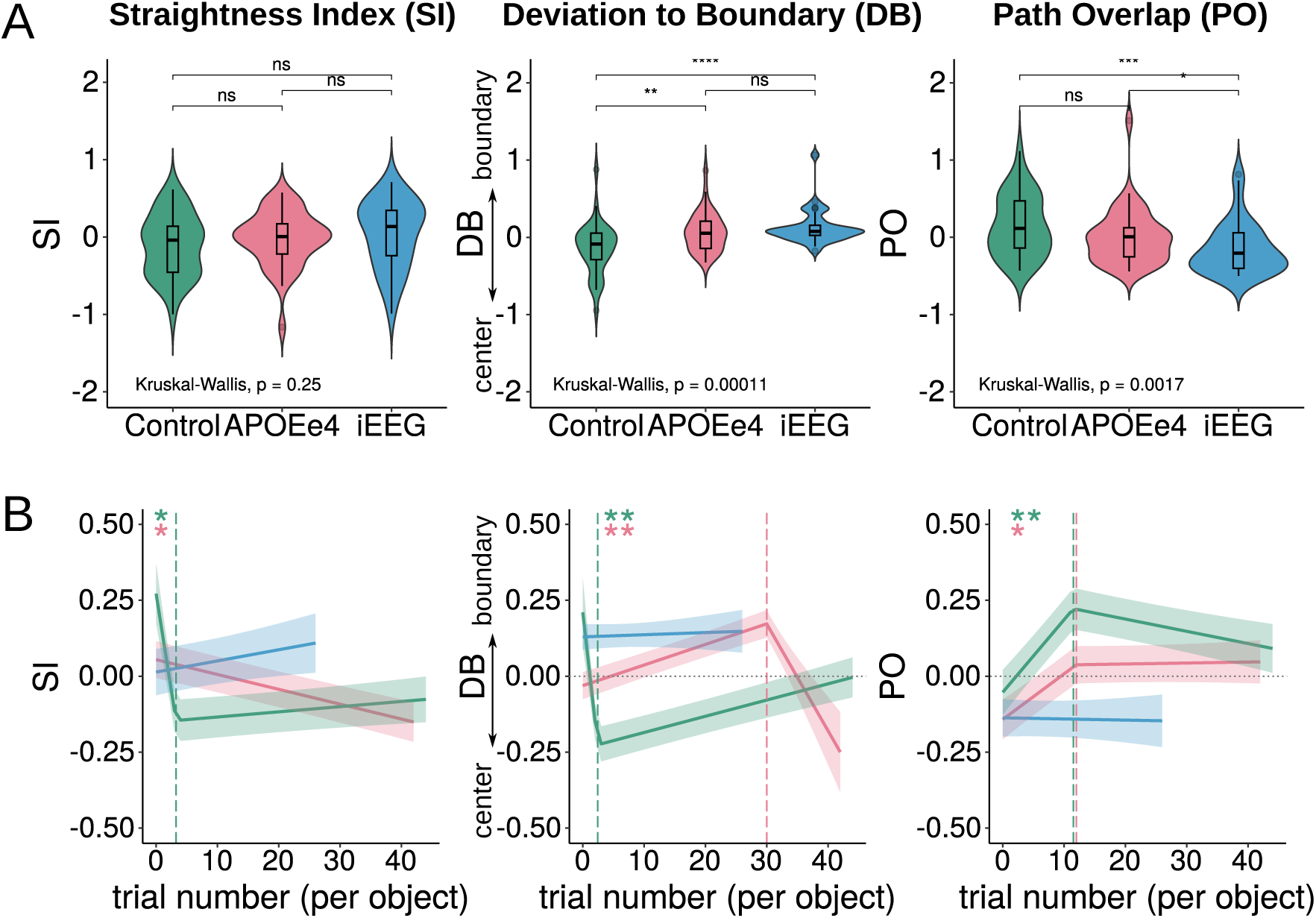
Group Differences in Strategy Use and Use of Strategies Changes Across the Experiment. A) Groups differed in their use of anchoring and route matching strategies. B) Controls and APOE-e4 groups showed differences in how they used strategies across the experiment. Significance (“*”) of the first slope (i.e. first leg of the piecewise regression) and breakpoint is shown after correcting for multiple comparisons using FDR (all p*<* 0.05). The significance of the second slope is also indicated by a “*” where applicable.

Supplementary analyses showed the location of the goal object, travel distance, speed and number of stops were all predictive of the metrics. For example, objects closer to the boundary resulted in straighter paths toward them, and similarly, when goal objects were further from the starting point (for full details see Supplementary Figure 3, Supplementary Table 6).

### Strategy Use Over the Course of Learning

Next, we investigated whether and how strategy use changed over the experiment. Significant changes in all three metrics were found in the Control and APOE-e4 groups (Figure 3B, Table 1, Supplemental Table 6), but not the iEEG group (thus iEEG results are not reported below). Controls took progressively less straight paths for the first 3 trials of each goal object (B=-1.3, p*≤* 0.001), with no significant change afterwards (B2=0.002, p*>*0.1). By contrast, the APOE-e4 group consistently went on less straight paths as the experiment progressed, with no significant breakpoint (B=-0.005, p*≤* 0.001). Controls initially (in the first 2-3 trials) deviated more and more towards the center of the arena (B=-1.8, p*≤* 0.001), after which this center-directed deviation diminished - but did not reverse into boundary-directed deviation; instead, overall deviation decreased (B = 0.005, p *≤.*001). Upon closer inspection, we found that when looking at the DB score for the first trial across all objects, Controls were significantly deviating towards the boundary (p*≤* 0.001), however already by the second trial (across all objects) this was not the case (p*>*0.1), and by the fourth trial they were significantly deviating towards the center of the arena (p=0.023) – thus the initial dip in DB score represents a rapid shift from deviating towards the boundary to deviating towards the center. The APOE4 group showed an inverted pattern, as they deviated increasingly towards the boundary until 30 trials per object, when they began to deviate less and less towards the boundary (B=0.007, p*≤* 0.001; B2=-0.035, p=0.001). To be exact, by the fourth trial the APOE-e4 group significantly deviated towards the boundary (p=0.002) and by the 32nd trial the deviation was no longer significant (p*>*0.1). Controls took increasingly overlapping paths in earlier trials and less overlapping paths in later trials (B= 0.024, p*≤* 0.001, B2=-0.004, p=0.03) while the APOE-e4 group took increasingly overlapping paths early in the experiment (B=0.015, p*≤* 0.001; B2=0.0003, p*>*0.1). These findings show that the overall group differences described above mainly reflected the patterns in early rather than late trials.

**Table 1:** Summary of Main Findings. For each strategy, whether it was adaptive overall as well as separately for early and late learning periods (relates to Figure 4), whether its use changed over the experiment (related to Figure 3B), and how it related to changes in hippocampal activity (related to Figure 6).

| Strategy | Group | Beneficial Strategy? | Change in use? | Hippocampal Relationship? |
| --- | --- | --- | --- | --- |
| Straightness (SI) | CONTROLS | Yes overall (late) | Decreasing early on then stable | Negative |
|  | APOE-e4 | Yes overall (early) | Decreasing over the course of learning | Negative |
|  | iEEG | Yes overall | No | Theta: Positive<br>Gamma: Negative |
| Deviation to Boundary (DB) | CONTROLS | Center: Yes overall (late),<br>Boundary: No | More deviation to center early on, then decreasing deviation | Center: Negative,<br>Boundary: Positive |
|  | APOE-e4 | Center: Yes overall (late),<br>Boundary: Yes overall (early) | More deviation to boundary for most of learning, then no deviation | Center: Negative,<br>Boundary: Positive |
|  | iEEG | No | No | Theta: Negative,<br>Gamma: Positive |
| Path Overlap (PO) | CONTROLS | Not overall (early) | Increasingly overlapping paths early on then reduction | Positive |
|  | APOE-e4 | No | No | Positive |
|  | iEEG | No | No | Theta: Negative,<br>Gamma: (Positive) |

### Strategy Use Predicts Performance

Having quantified the three path metrics for every trial, we tested whether they predicted performance on the task, i.e. trial-wise drop error, again using piecewise regression analyses with the different strategy metrics as predictors (separately), and (corrected) drop error as dependent variable (Figure 4A; see also Table 1). In Controls, we found that going on more straight paths (higher SI) predicted better performance (lower drop error) until a breakpoint (B=-0.59, p*≤* 0.001), after which the relationship reversed and became significant in the opposite direction (B2=14.94, p*≤* 0.001). Notably, this reversal occurred for very straight paths, and putatively reflects instances where participants were unsure and travelled in a short straight line to terminate the trial (see Supplemental Figure 4 for details). Deviating towards the center (DB*<*0) was related to better performance (B=0.75, p*≤* 0.001), while deviating towards the boundary was not (B2=-0.197, p*>* 0.1) – in line with the overall tendency of Controls to deviate to the center (see also Kunz et al 2015). Another preferred strategy of Controls, travelling along overlapping routes, was also beneficial until a certain point when the benefit plateaued (B= −1.45, p*≤* 0.001; B2= −0.063, p*>*0.1) and this point was at about 3% increase in path overlap.

**Figure 4:**
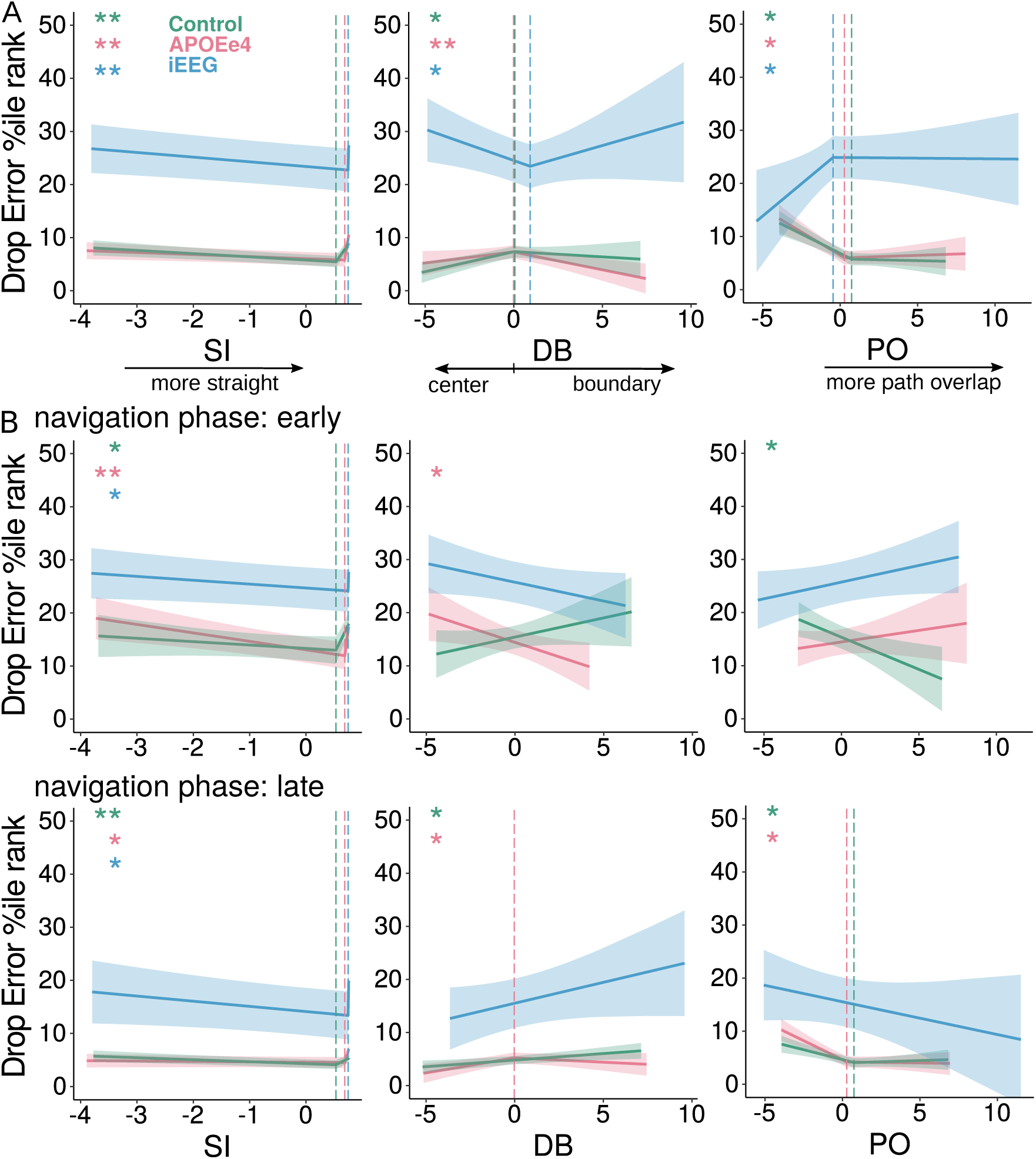
Performance is Related to Strategy Use. A) Relationship of path metrics and performance across the entire experiment. B) Relationship of path metrics and performance separately for the early and late learning phases, as seen in Figure 1D. Significance (“*”) of the first slope (i.e. first leg of the piecewise regression) and breakpoint is shown after correcting for multiple comparisons using FDR (all p*<*0.05). The significance of the second slope is also indicated by a “*” where applicable (note: second leg only marked as significant if the breakpoint and second leg were significant). If there is no breakpoint (i.e. no dashed line), then a linear model was fitted as a piecewise model provided no additional benefit.

The APOE-e4 group showed a similar pattern for both straightness (B=-0.38, p=0.008; B2=67.79, p*≤* 0.001) and overlapping paths (B=-1.75, p*≤* 0.001; B2=0.097, p*>*0.1). Contrary to Controls, however, this group benefitted from deviating towards the center and the boundary the arena (B1=0.42, p=0.044, B2= −0.67, p*≤* 0.001). In the iEEG group, only straightness was associated with better performance, as in the other groups (B=-0.88, p=0.013; B2=427.32, p*≤* 0.001). No clear benefit of the other two strategies was found, in fact deviating to the center and increasingly overlapping paths were detrimental to performance. Potentially, this could be a result of the large variability in performance in this group thus obscuring any clear beneficial use of strategies.

For completeness, we also examined the results of the piecewise regressions when including the other metrics as covariates in the model (e.g.: predictor variables would be main effect of Straightness and the corresponding interaction term, as well as main effects of Deviation to Boundary and Overlap of Paths). We found the results were qualitatively very similar (see Supplemental Table 7B).

We also investigated, given the changes in strategy use (Figure 3B) and reductions in Drop Error (Figure 1D) over the course of the experiment, the relationship between strategies and Drop Error separately for the early phase (up to 6 trials per object in the Control/APOE-e4 groups, and 9 trials per object in the iEEG group; see Figure 1D), and the late phase (all trials after the early phase) separately (thus breakpoints were the same as above, but only early - or late - learning phase trials were included in each model). During the early phase of learning, Controls benefitted from taking increasingly overlapping paths (early PO: B= −1.21, p=0.009), and APOE -e4 carriers benefitted from deviating towards the boundary (B= −1.09, p=0.025), and going on straighter paths until the breakpoint (B=-1.59, p=0.001; B2=77.51, p*≤* 0.001) (Figure 4B top panel). In later learning, Controls benefitted from going on straighter paths until the breakpoint (B= −0.37, p=0.002; B2=5.38 p*≤* 0.001) and taking overlapping paths up until a point (B=-0.73, p*≤* 0.001; B2=0.09, p*>*0.1), while the APOE-e4 group showed a benefit only from deviating towards the center of the arena (B=0.57, p=0.001, B2=-0.16, p*>*0.1) rather than the boundary as early in learning. They also showed a similar pattern to controls in the benefit of using overlapping paths (B=-1.42, p*≤* 0.001; B2=0.04, p*>*0.1, Figure 4B lower panel). No effects were observed for the iEEG group.

In summary, these analyses again show that the overall preference of the different groups (deviation to the boundary in APOE-e4 carriers and using an egocentric strategy in Controls) corresponded to the benefit in earlier rather than later trials. They also suggest that in Controls, switching from a more egocentric strategy early in learning (route matching) to putative cognitive map use (straight trials) during later learning stages benefits performance. The APOE-e4 carriers, however, consistently use the geometry of the arena, first adaptively using the boundary and then adaptively deviating towards the center.

### Hippocampal Recruitment Benefits Performance only in APOe4 carriers and iEEG patients

We investigated if hippocampal activity was associated with better performance. Hippocampal activity was measured by extracting bilateral (see Methods for details) BOLD activity in Control and APOE-e4 participants, and theta power (3-8Hz) and high gamma power (60-90Hz) in iEEG patients (electrode locations plotted in Supplemental Figure 10), timepoint-by-timepoint. Mixed-effects models were corrected for temporal autocorrelations of neural activity, see Methods.

In Controls, there was no overall relationship between hippocampal BOLD activity and drop error (p*>*0.1). By contrast, in the APOE-e4 group higher hippocampal BOLD was associated with lower drop errors (B= −0.0009, p=0.001). In the iEEG group, theta power (3-8Hz averaged) was unrelated to performance (p*>*0.1), but higher gamma (60-90Hz) power predicted better performance (B= −0.00015, p=0.001) (Figure 5A).

**Figure 5:**
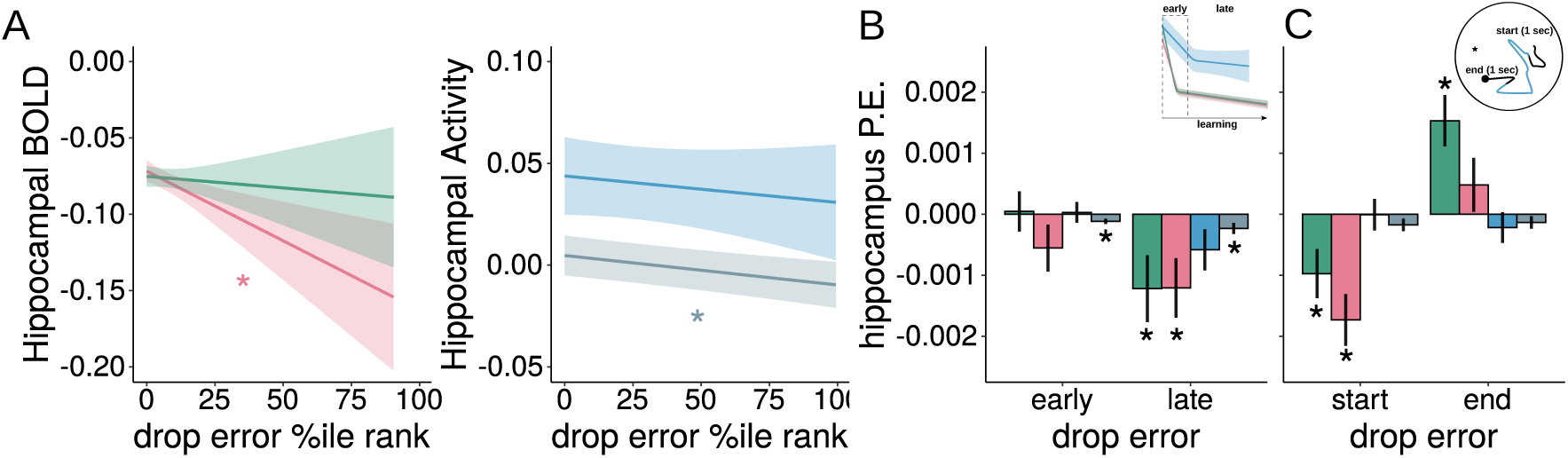
The hippocampus is recruited to aid performance in group-specific manner A) Performance, i.e. corrected Drop Error, was used to predict hippocampal activity. Left: Hippocampal BOLD activity predicted better performance in APOE-e4 carriers, but not in Controls. Right: In the iEEG group, higher gamma power was related to better performance (gray line), while no effect was found for theta activity (blue line). B) Relationships of Drop Error and hippocampal activity (extracted parameter estimates plotted) separately for early vs late learning phases and C) for the first (”start”) and last (”end”) second of every trial. Plotted are the extracted beta estimates for each phase/trial period, per group. Results are corrected for FDR (* p*<* 0.05).

When looking at early and late learning phases separately, BOLD activity in both Controls and APOE-e4 carriers was associated with better performance during late learning (both B=-0.001, both p*<*0.023), but not during early learning (p*>*.01). By contrast, in the iEEG group, gamma power consistently predicted performance both during the early (B= −0.0001, p=0.021) and the late learning period (B= −0.0002, p=0.042) (Figure 5B left).

An alternative way to understand how hippocampal activity relates to performance is to analyze activity at the start (first second during retrieval) or end (last second during retrieval right before drop) of a trial (Figure 5C right inset). We found a benefit of hippocampal recruitment during the initial part of the navigation trajectory in both Controls and APOE-e4 carriers (Figure 5C, Controls B= −0.001, p=0.016; APOE-e4 B= −0.002, p=0.001), indicating some form of beneficial retrieval-related activity as to the intended goal location. Interestingly, in Controls we also found a significant effect at the end of the retrieval trial (i.e. last second before drop) – but in the opposite direction, thus more hippocampal BOLD reflected higher Drop Error (B=0.002, p=0.001). This might relate to more extensive learning in trials with high drop error, in which Controls may engage in more re-encoding (supported by the fact that hippocampal activity in both Control and APOE-e4 groups is overall higher in the reencoding phase than during retrieval, Supplemental Figure 5). Previous studies suggest that entorhinal cortex could be dysfunctional even at early ages in APOE-e4 carriers [Kunz et al., 2015] and that in addition to the MTL, striatal networks contribute to navigation as well [Bohbot et al., 2007, Goodroe et al., 2018, Henke, 2010]. We thus investigated the relationship between drop error and neural activity in bilateral entorhinal cortex and bilateral caudate nucleus in the Control and APOE-e4 group (for details on ROIs and for details on why the entorhinal contacts in the iEEG group were not used in this analysis please see Methods-Preprocessing). We did not find that drop error was significantly related to neural activity in bilateral entorhinal cortex in either group (p*>*0.1). However, increased BOLD in the caudate was related to better performance in the APOE-e4 group (B=-0.001, p=0.030, Supplemental Figure 9 and Supplemental Table 13).

### Hippocampal Recruitment is Dependent on Adaptive Strategy Use and is Group Specific

While all strategies seem to be adaptive (i.e., beneficial for performance) to some extent depending on group membership, we next investigated their neural underpinnings. In addition, we aimed to understand the relationship between a particular strategy use and neural activity as a function of whether it was adaptive or not. We therefore used the breakpoints from the Drop Error analysis (and associated path metric interaction terms) in mixed-effect models to predict hippocampal activity (Figure 6).

**Figure 6:**
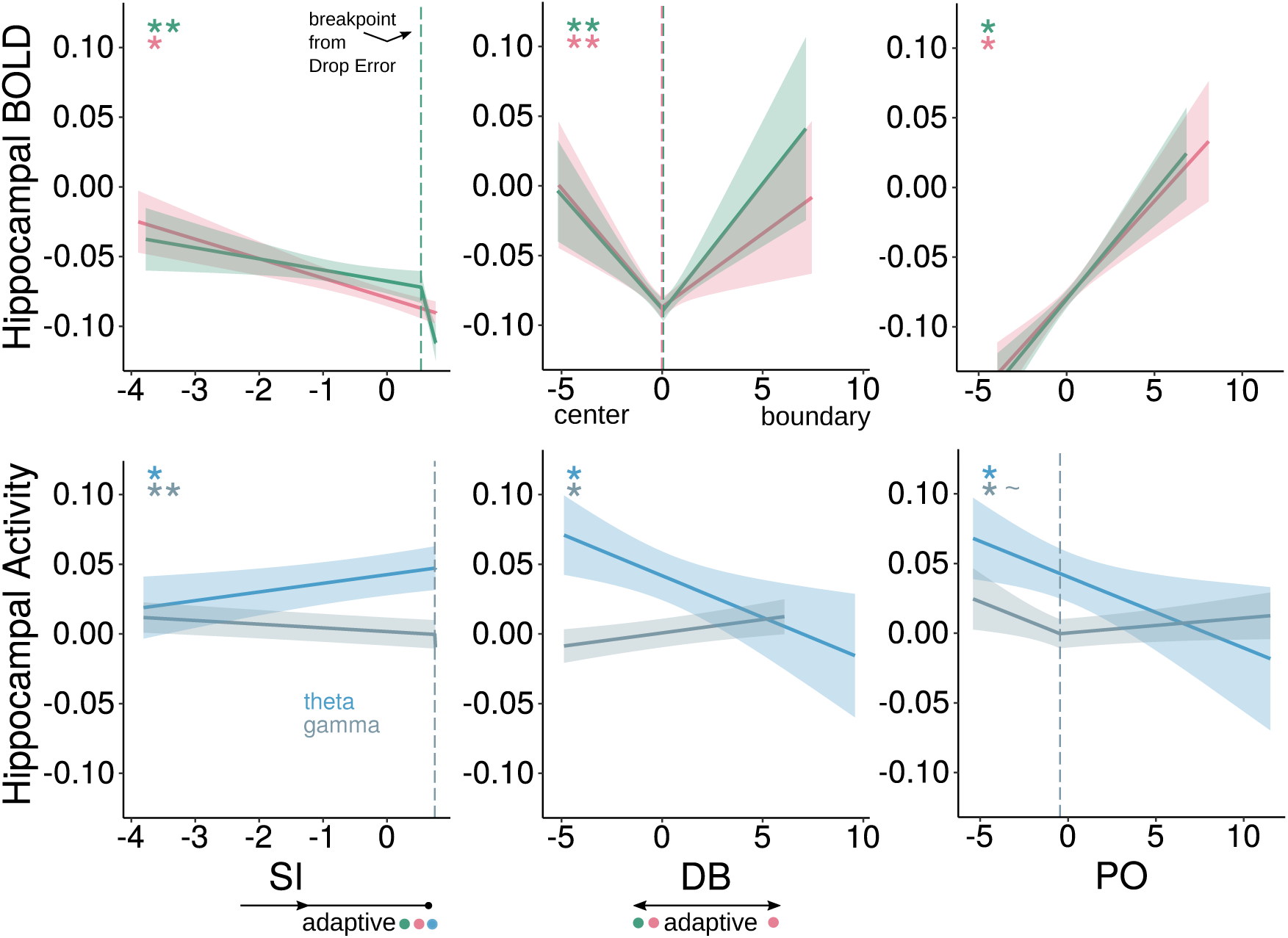
Hippocampus Is Involved Differently Depending on Adaptive Strategy Use. We used piecewise mixed-effects models, with breakpoints taken as established for Drop Error (Figure 4), in order to understand how hippocampal activity related to strategy use as a function of whether it was beneficial or not. In Controls and APOE-e4, more straight paths (SI), leading to better performance, were associated with lower hippocampal BOLD activity, while deviating either to the center or the boundary was associated with higher hippocampal BOLD activity, which was also related to better performance (albeit adaptive deviation to either center or boundary was only present in APOE-e4s and only adaptive deviation to center in Controls). This implies that changes in the direction of BOLD response in the hippocampus can occur as a function of the type of adaptive strategy use. Hippocampal BOLD also increased when taking more overlapping paths, although this was not clearly related to performance. In the iEEG group, higher theta power and lower gamma power were associated with adaptively taking straighter paths, theta power decreased when deviating more towards the boundary and route matching, and gamma activity increased when deviating towards the boundary. Significance (“*”) is shown after correcting for multiple comparisons using FDR.

In Controls, going on straighter paths (which led to better performance) was associated with decreased BOLD activity in the hippocampus; for very straight paths that were associated with poor performance, this decrease in BOLD was even steeper (B1= −0.008, p=0.024, B2= −0.17, p*≤* 0.001). Adaptive deviation towards the center of the arena was associated with increased BOLD activity, as was deviation towards the boundary, despite the latter not relating to performance in the Controls (B1= −0.016, p=0.001, B2= 0.018, p*≤* 0.001). Taking more overlapping paths (route matching), while beneficial to a point, was associated with a linear increase in hippocampal BOLD (B= 0.015, p=0.001). These findings show that in Controls some adaptive strategies were associated with reductions and others with increases of hippocampal BOLD activity, which may explain the lack of an overall linear relationship between BOLD activity and performance in this group (Figure 5A).

In the APOE-e4 carrier group, going on straighter paths was associated with a decrease of hippocampal BOLD activity, independent of whether it was adaptive or not (B=-0.014, p=0.001). Adaptive deviation towards the center and boundary of the arena was associated with increased BOLD activity (B1= −0.017, p=0.001; B2=0.011, p=0.008). As in Controls, taking overlapping paths (route matching) was associated with higher hippocampal BOLD (B=0.014, p=0.001). Thus, “anchoring” at the boundary – the predominant strategy in this group – was associated with higher hippocampal BOLD activity, putatively accounting for the overall benefit of hippocampal recruitment.

In the iEEG group, going on straighter paths was associated with a significant linear increase in hippocampal theta power (and decrease in gamma power; theta: B=0.006, p=0.014; gamma: B=-0.003, p=0.001). Deviating towards the boundary and taking increasingly overlapping paths led to significant linear decreases in theta power (DB: B= −0.006, p=0.012; PO: B=-0.005, p=0.018), and deviating towards the boundary was in addition associated with significantly increased gamma power (B=0.002, p=0.007). See also Supplemental Figure 6 and Table 12 for results at individual theta frequencies. Thus, anchoring and route matching strategies were associated with lower theta power, while straighter paths were associated with increased theta power – and theta power overall was inversely related to BOLD activity. Increased gamma power was overall beneficial to performance (Figure 5A right) and was also increased when deviating to boundary (only), which is the strategy that iEEG patients used more than the other groups, albeit without being beneficial to them. However, gamma power was also increased for low SI/PO values (Figure 6 lower left/right), i.e. when patients were taking non-matching routes, putatively when they were moving on very idiosyncratic paths. Interestingly, this was also when they showed significant decrease in Drop Error (Figure 4A right), thus the overall performance benefit of higher gamma power in this group may be driven by these curvy non-typical paths. It is worth noting that while behaviorally very straight trials did relate positively to speed (Supplemental Figure 3), we did not find that higher speeds resulted in higher theta power (Supplemental Figure 8C) and thus our finding of higher theta power with increasingly straight paths is unlikely to be the result of speed changes.

In summary, while overall, there was an inverse relationship between BOLD activity and theta power (vs gamma power), adaptive strategy use was not always associated with increases or decreases in hippocampal involvement, but rather hippocampal activity was strategy and group specific. Beneficial strategies in the Control and APOE-e4 groups, such as using (modestly) straight paths and deviating to the center/boundary, were associated with opposite (lower and higher, respectively) BOLD activities, and route matching was associated with increased BOLD, despite not conferring any additional benefit after a certain point. These results suggest that changes in direction of BOLD activity are not directly reflecting a specific strategy but also indicate whether it is adaptive or not (see Discussion).

We also investigated how the relationship between strategies and hippocampal activity changes as a function of learning (early vs late) and trial period (start vs end), see Supplemental Figure 7 and Supplemental Table 10 & 11. For completeness, we also report results of predictors for hippocampal power consisting of behavioral variables similar to those relating to previous reports (e.g.: distance to drop location, speed etc., see Supplemental Figure 8), as well as the relationship between strategies and hippocampal theta power per individual frequency (i.e. 3-8Hz separately, Supplemental Figure 6).

## Discussion

Navigation behavior is complex: idiosyncratic [Wolbers and Hegarty, 2010], dependent on context and environment[Coutrot et al., 2018, Ekstrom and Isham, 2017], and changing with age [Bohbot et al., 2012, Bullens et al., 2010, West et al., 2023] and disease [Segen et al., 2022]. Understanding how and under what conditions different strategies are employed to solve navigation tasks is still debated [Hegarty et al., 2023]. One approach is to design experimental paradigms that include different conditions to emphasize the use of one strategy over another (hypothesis-driven and externally controlled, e.g.: [Nett et al., 2025]). An alternative would be to use a completely data driven approach, such as in [Donnarumma et al., 2021], A blend of the two approaches, hypothesis-and data-driven, is what we considered here, inspired by the idea that changes in neural coding can often be explained by covert processes related to changing internal cognitive variables [Johnson et al., 2009], and by the importance of understanding how and under what circumstances certain behaviors are elicited [Krakauer et al., 2017].

We found that navigation strategy was not uniquely a trait-level characteristic. Instead, it was informative to quantify navigation trajectories across trials (Figure 2B) and across multiple dimensions simultaneously, an approach that fits with previous frameworks in which egocentric and allocentric distinctions are not mutually exclusive or clearly indicative of a lack or presence of a Euclidean ‘cognitive map’ [Filimon, 2015, Peer et al., 2021, Warren et al., 2017]. We also found that our metrics were specific to active goal-directed navigation (retrieval phase) not general movement without spatial memory component (reencoding phase, see Supplemental Figure 2A), and that their use shifted over the course of the experiment (Figure 3B).

### Going on a straighter path leads to better performance

We hypothesized that more direct routes towards a goal location, irrespective of the start location, would be indicative of the presence of some form of “cognitive map” – putatively an allocentric representation. We found that going on straighter paths was adaptive for all groups (Figure 4A left), i.e. resulted in better spatial memory performance. Interestingly, Controls only engaged in straight paths as learning progressed and only benefitted from straight paths in the late learning period, while APOE-e4 carriers took less and less straight paths as learning progressed and only benefitted from this strategy early in learning. The iEEG group showed an overall benefit of straight paths. In APOE-e4 carriers, going on straighter paths always related to reduced hippocampal BOLD activity (Figure 6, Supplemental Figure 7 top left); in Controls, while the pattern was similar, there was an additional reduction in hippocampal BOLD for the very straight nonadaptive trials (Figure 6, second leg), as well as in the caudate (Supplementary Figure 9). This may be a signal in Controls that there is uncertainty and straight movement is not adaptive. In the iEEG group, we found that straighter paths resulted in increased theta and decreased gamma power in the hippocampus, with the latter band mirroring the increased reduction for nonadaptive trials as seen in Controls.

### Anchoring to the center or boundary of arena differentiate Controls from APOE-e4 carriers and relates to different neural mechanisms

As predicted from previous reports [Bierbrauer et al., 2020, Coughlan et al., 2019, Kunz et al., 2015], APOE-e4 carriers used the boundary of the arena to their advantage. However, when dividing learning into phases, we found deviating to the boundary was adaptive in the early phase, but deviating to the center was adaptive in the later phase (Figure 4B). By contrast, Controls only benefitted from deviating to the center. Thus, APOE-e4 carriers behave like Control participants after initial learning has taken place.

Deviating to the center may provide a vantage point to observe the relationship between distant landmarks to estimate goal directions as well as distances (whereas the boundary can provide distance, but not direction information). Interestingly, in both groups, deviation to both the center of the arena and the boundary was related to increased BOLD activity in hippocampus (Figure 6 top middle) and entorhinal cortex (Supplemental Figure 9 top). Furthermore, Controls showed an increase in BOLD activity when deviating towards the boundary in the caudate nucleus (Supplemental Figure 9 bottom), suggesting that increased BOLD when deviating towards the boundary may relate to response learning mechanisms. While the iEEG group did not benefit from deviating to either center or boundary, increased deviation to the boundary was associated with decreased theta and increased gamma power (Figure 6 bottom middle).

Speculatively, the deviation of APOE-e4 carriers towards the boundary reflects a compensatory behavior in which they shift from using boundaries as allocentric cues to egocentric anchoring points, perhaps in order to gauge distances [Hartley et al., 2004]. In previous studies, APOE-e4 carriers strongly relied on local landmarks [Bierbrauer et al., 2020]; but see [Lim et al., 2023]; similar to a landmark, a boundary can be used to estimate distances, and in combination with distal cues can be used to triangulate goals. Furthermore, while the boundary vector hypothesis [Barry et al., 2006, Bicanski and Burgess, 2018, Lever et al., 2009] emphasizes the role of cells coding direction and distance to boundaries as providing the input for allocentric representations such as place cells, boundaries have been shown to provide egocentric information both in rodents [Gofman et al., 2019] and in the form of egocentric bearing cells in humans [Kunz et al., 2021]. Thus, neural reference points within the medial temporal lobe to the boundary or behaviorally manifested boundary-hugging could be a useful approach for building a representation of the available space. Equally, there has been evidence that cells representing the direction towards the boundaries are overrepresented in the center of the arena space [LaChance et al., 2019, Wang et al., 2018], which would explain why in Controls deviating towards the center is useful, and later in navigation, also for the APOE-e4 group. How the strategy of deviation towards the boundary is related to changes in egocentric coding of boundaries - which has been proposed to be instrumental in the transformation or generation of allocentric codes [Alexander et al., 2023, Bicanski and Burgess, 2020] - remains an open question.

### Taking the same route to the goal is not useful but it activates medial temporal areas

Route-matching is considered an egocentric strategy as it relies on a self-referenced set of actions in relation to landmarks or viewpoints in the environment. Use of heading directions (viewpoint matching) has been reported previously to differentiate strategy use in the radial maze [Igĺoi et al., 2009], where rotating to look around at the start of an arm resulted in allocentric strategy use, whereas rotations in the center were related to correcting an initially egocentric strategy to an allocentric strategy for completing the maze. Here we considered this strategy by quantifying the overlap between routes taken to a particular goal location. This strategy was employed by and adaptive for Controls early in learning (Figure 3B right and Figure 4B top right). Previous research has found that when not forced to, people will retrace a longer known path rather than take a shortcut [Marchette et al., 2011]. Our definition of route (and by definition, viewpoint) matching is reminiscent of sequential-egocentric navigation during which goals are reached via visiting a sequence of subgoals [Nyberg et al., 2022, Shamash et al., 2021]. Interestingly, overlapping paths were related to increased BOLD activity in hippocampus and entorhinal cortex, which may be considered surprising as viewpoint dependence has been reported in a patient with severe hippocampal damage [King et al., 2002]. The increase in BOLD response may simply reflect the recall of previously taken routes, in line with theories placing the medial temporal lobe as the core area supporting episodic retrieval, i.e. the binding of items in context [Moscovitch et al., 2005, Moscovitch et al., 2006, Ranganath, 2010] or enabling the recall of a sequence of events from memory as the basis for the ability to navigate through a sequence of physical spaces [Eichenbaum, 2017].

### Meso- to Macroscale: not simply higher BOLD, lower theta power

Generally, we find a positive relationship between high frequency gamma power measured in iEEG recordings and BOLD activity measured with fMRI, and an inverse relationship between BOLD activity and theta oscillations (Table 1). These findings are in line with previous work showing that low-frequency oscillations are negatively correlated with BOLD activity while the opposite is the case for higher frequency activity in the gamma frequency range (for reviews see: [Ekstrom, 2010, Kunz et al., 2019a]). Only one study so far has been able to show a positive relationship between theta power and BOLD activity, but this was found in parahippocampal cortex, not the hippocampus [Ekstrom et al., 2008]. However, we also found that deviation to the center was associated with increases in both hippocampal theta power and BOLD activity (Figure 6, middle panels). This might be explained by recent findings that metabolic activity is not consistently antagonistic when comparing positive and negative BOLD changes [**?**, Stiernman et al., 2021], and the hippocampus exhibits functionally different neurovascular coupling compared to neocortex [Shaw et al., 2021], with some research showing that high amplitude BOLD events may be accompanied by decreases in LFP power [Zhang et al., 2020].

### Additional recruitment of neural resources in APOE e4 carriers

Independently of strategy use, we found that APOE-e4 carriers may have been able to perform as well as Controls using compensatory mechanisms, in the form of recruiting hippocampus and caudate (Figure 5 and Supplemental Figure 9). This effect was absent in Controls, which dovetails with previous findings of increased medial temporal activation in 4 carriers [Filippini et al., 2009]. This may also be related to findings that the hippocampus shows altered neurovascular coupling in APOE-e4 carrier mice already at early ages, including reduced vascular responsiveness and vasomotion, which may contribute to their increased risk of A deposition later in life [Bonnar et al., 2023]. Additionally, the common finding that cerebral blood flow is increased in young healthy APOE-e4 carriers has been shown to be accompanied by sub-optimal tissue oxygen uptake (microvascular dysfunction, [Aamand et al., 2024]), therefore changes in the recruitment of medial temporal areas during our spatial memory task could be related to these microvascular changes.

### From spatial memory to planning to navigation: strategies are just part of the story

Studies in humans have established the presence of theta-gamma coupling and its functional role in memory encoding and retrieval [Heusser et al., 2016, Lega et al., 2012, Lega et al., 2016], and more recently, how goals are represented at distinct theta phases [Kunz et al., 2019b]. How these relationships may be modulated by strategies taken to the goal remains an open question. However, we did find that hippocampal BOLD activity changed across the retrieval phase of navigation in a strategy- and group specific manner (early vs late & start vs end of trial, Supplementary Figure 7), thus we can speculate that goals are represented differently depending on how a route is planned, putatively relating to whether the prefrontal cortex is involved [Patai and Spiers, 2021]. Furthermore, it remains to be investigated whether and how the changes in strategy use we observed in our study (e.g. Controls shift from deviating to center to going on straighter paths) relate to transformation of reference frames (egocentric to allocentric) across learning, putatively via the retrosplenial cortex [Alexander et al., 2023, Epstein, 2008].

## Conclusion

Our results show that navigation behavior even in a simple circular arena is highly variable, both within individuals and across learning, and is best described as a combination of parallel strategies. External variables such as goal locations also influence the deployment of strategies, underscoring the need to understand the conditions and contexts that may lead to changes in how navigation problems are solved. Adding to our understanding of early diagnosis for dementia risk, our results highlight that APOE-e4 carriers are not just using different strategies from Controls, but are adaptive and changing, indicating that time-resolved and/or longitudinal studies are needed to map maladaptive behavior (and neural mechanisms) that may indicate susceptibility to disease. We lay a foundation to use these features to classify individuals and their instantaneous behavioral states and contribute to areas of research aiming to understand the relationships between meso-and macroscale neural recordings in the context of ongoing volitional behavior.

## Methods

### Participants

We analyzed pre-existing datasets from three groups of participants: fMRI data from 37 Controls (APOE e3/e3) and 38 APOE-e4 carriers (APOE e3/e4) as published in [Kunz et al., 2015]. iEEG data was collected from 35 patients with pharmaco-refractory epilepsy (some patients included in [Kunz et al., 2019b, Kunz et al., 2024]), over 47 sessions (some participants performed multiple sessions over multiple days), and only participants with hippocampal contacts were included in this analysis. One Control participant performed the task by taking paths that always touched the boundary in the same south-east corner of the arena (on all trials, to all goal locations). While this behavior is strategic and interesting, we have opted to exclude them from further analysis (this participant showed a Deviation to Boundary (DB) that was 4 times larger than the interquartile range).

Our final dataset thus consisted of 36 Controls (18 females, mean age = 22.83 (18-30)), 38 APOE-e4 carriers (20 females, mean age = 22.34 (18-29)), and 33 intracranial patients across 45 sessions (19 females, mean age = 35.3 (19-61)).

### Navigation task

Participants performed an object-location memory task in which they navigated freely between hidden goal locations in a circular virtual arena, with distal landmarks located behind a circular wall. Participants were asked to navigate toward and memorize the locations of 8 different everyday objects (location placement was selected from 16 possible goal locations for the fMRI groups, and individually determined from a larger set of locations for the iEEG group, but in all cases the distribution of objects relative to the boundary was matched across participants/groups). Each trial consisted of a cue presentation phase, a retrieval phase, a feedback phase, and a re-encoding phase (Figure 1A). The duration of the retrieval phase was self-paced. When participants reached the location they considered correct, they “dropped” the object by hitting the space bar (iEEG group) or the backward arrow key (Controls and APOE-e4 group). The object then appeared at its correct location, and patients navigated to this location (‘reencoding phase’). The next trial (cue for the next object) started from this location, i.e. the just encoded goal location. For details please see description in [Kunz et al., 2019b] for iEEG version and [Kunz et al., 2015] for fMRI version. Participants completed variable numbers of trials. Controls: trials performed = 230.59 (107-323), minimum/maximum times each object was visited: 11/44. APOE-e4: trials performed = 230.68 (149-320), minimum/maximum times each object was visited: 17/42. iEEG: trials performed = 130.72 (15-324), minimum/maximum times each object was visited: 2-46.

For the analysis presented in this paper, we only look at the retrieval phase (and reencoding phase as a control period, see Supplemental Figure 2) to extract navigation paths to the goal locations, see Path Indices section below.

### Preprocessing

For the fMRI dataset, we used preprocessed data (slice-time correction, reslicing, smoothing), please see [Kunz et al., 2015] for details. In brief, BOLD activity was modelled across the entire dataset using the standard six motion nuisance regressors (using SPM) and accounting for temporal autocorrelation (AR(1)). The beta values for each volume (i.e. each TR of 2.6sec) were extracted from the bilateral hippocampus and entorhinal cortex (native space, Freesurfer segmentation, labels: Right-Hippocampus Left-Hippocampus, ctx-rh-entorhinal and ctx-lh-entorhinal). Neural activity across the entire retrieval period was subsequently related to behavioral metrics (see Path Indices section below).

The iEEG dataset consisted of a sample of recordings already reported [Kunz et al., 2019b], as well as new data collected from the Department of Epileptology, University of Freiburg, Freiburg im Breisgau, Germany. Patients underwent a surgical procedure in which electrodes (Ad-Tech, Racine, WI, USA) were implanted stereotactically deep within the brain parenchyma. Electrode placements were determined solely on the basis of clinical considerations. Each patient had a unique implantation scheme with a unique anatomical distribution of electrode channels in various brain regions. iEEG data were acquired using a Compumedics system (Compumedics, Abbotsford, Victoria, Australia) at a sampling rate of 2000Hz. Signals were referenced to Cz.

Time-periods with interictal seizure activity were excluded manually after visual inspection. Data from each electrode contact was re-referenced to a bipolar montage, high-pass filtered with a lower bound of 0.2Hz, and activity higher than 5 times the interquartile range was labelled as artifacts. Wavelet convolution (Morlet) was performed with the MNE package https://zenodo.org/records/10161630 between 3-8Hz/60-90Hz (evenly spaced on logarithmic scale), with 5 cycles per frequency. Time-frequency data was then log transformed per frequency per timepoint, and finally z-transformed, thus we obtained what we label as ‘theta’ or ‘gamma’ power for all retrieval period timepoints across each dataset. Data was then epoched to match the sampling of the behavioral data (0.1 sec) and aligned to the behavioral data in order to be related to behavioral metrics (see Path Indices section below). Electrode localization (on bipolar virtual electrodes) was performed using FSL https://fsl.fmrib.ox.ac.uk/fsl/fslwiki/FSL and PyLocator http://pylocator.thorstenkranz.de/. Channel locations were manually identified and for this analysis, we focus on hippocampal electrodes only. Data from bilateral hippocampal electrodes was averaged for analysis. Total electrodes used: 111 (58 from right, across 24 patients; 53 from left hippocampus, across 19 patients, Supplemental Figure 10). We did not explore entorhinal activity in the iEEG sample as only 8 patients had contact in this region.

### Performance on the Task

Drop Error was corrected for the fact that by chance, error would be lower for goal object locations located in the center of the arena (i.e. max error would be the radius of the arena). We followed the procedure reported in [Kunz et al., 2021, Kunz et al., 2024]. A sample of random drop locations was generated (n=1000) and the Drop Error for all of these random locations was calculated. These were then compared to the actual observed Drop Error on each trial, by using a percentile ranking score which gave the percentage of random drops below of a given observed Drop Error (“Drop Error %ile rank” in the figures). Thus, if the actual Drop Error was very low, say the participant was very accurate, then the percentile rank of the Drop Error would also be very low, because only a low percentage of random drops would be as good as (or better than) the observed Drop Error.

### Path Indices: Straightness, Deviation to Boundary, Path Overlap

We aimed to quantify putative strategies by using a hybrid hypothesis- and data-driven approach to characterize paths taken during navigation by participants as they located the hidden objects in the arena. To this end we defined four metrics: straightness of the paths taken, amount the path taken deviated towards the boundary of the arena, the amount of path overlap between trials to a particular goal location, and a similar metric, the amount of heading direction overlap between trials to a particular goal location. All metrics were calculated in relation to the ‘optimal’ or direct path between the start location (which was always the actual goal location from the previous trial, reached in the reencoding phase) and the final drop location (which is not necessarily the goal location, but the location indicated by the participant, where they believed the object to be, i.e. ‘drop location’). Using the optimal paths was important to establish a ‘ground truth’ to compare the paths taken by the participant, as the paths were by definition constrained by the start locations as well as the boundary of the arena. We will use the term ‘observed’ path to refer to the path actually taken by the participant on a given trial. The calculation of the metrics in detail were as follows:

1. Straightness Index: the ratio between the length of the direct path and the observed path taken from the start location to the drop location. The direct path was calculated as the Euclidian distance between the x/y coordinates of the start and drop location. The observed path was calculated as the total length travelled based on paired (Euclidian) distances between subsequent x/y locations in the trial. A value close to 1 would indicate a straighter path taken by the participant. This metric is measured trial-by-trial. We consider this a measure of allocentric strategies, as approaching variable goal locations from variable start locations implies an understanding of the relations between locations and distal landmarks.
2. Deviation to the Boundary: first we measured the distance, for each x/y location in a path (optimal and observed), from the boundary (operationalised as Euclidian distance from the centre minus the arena radius). Note for the optimal path, we created a vector between the start and drop location, using evenly spaced samples, based on the size of the observed path. Then at each point in the path, we took the difference between the current distance to the boundary for the optimal and the observed path. A positive value would indicate that the observed path deviated towards the boundary, a negative that the observed path deviated towards the center. This metric can be measured time-point by time-point and can also be measured trial-by-trial by taking the median deviation values across the path (henceforth shortened to as DB). We consider this a measure of an anchoring strategy, in which the boundary of the arena may be used as a form of landmark, to provide distance information.
3. Path Overlap (PO): the amount that taken paths overlap towards a given goal location. This is calculated by taking any given trial i to goal object x, and comparing it to all other trials (except i) that were taken to that particular goal location, using percentage of x/y coordinates overlapping. In order to quantify the percentage overlap, paths were first downsampled (into a 2D histogram of 50×50 pixels (from 5000×5000 of the arena), binarised and then convolved, i.e. smoothed, by factor of 2. Then we performed a bitwise operation to establish the number of matching and total pixels in the images, and percentage was calculated from 100*matches/total pixels. The percentage overlap with all trials to the goal object was calculated for both the observed path and the optimal path for trial i. The difference between the observed path overlap percentage (trial i) and direct path overlap percentage (trial i) gives a trial-by-trial measure to establish how stereotypical each path was when approaching a specific goal location. We consider this a measure of a type of egocentric strategy, ie stereotypicality in movement paths, or route-matching.

### Z-scoring

Z-scoring was performed across all data points for relevant behavioral variables, with all three participant groups combined. However, as some metrics varied only at the trial level (eg: Path metrics: SI/DB/PO), while others were time-point by time-point measures (eg: Distance to Drop Location, Deviation to Boundary), for trial-level variables, we calculated the mean and standard deviation for z-scoring based on the trial-averaged data, to ensure that the length of the trial (in samples), with a given metric repeated for the entire trial, did not skew the descriptive statistics. We opted for z-scores as this facilitated comparison across path indices, which were on different scale to begin with (Straightness is bounded *ɛ* to 1, whereas Deviation to Boundary theoretically can be −5000 to +5000, which is the size of the arena radius). Additionally for easier interpretation of mixed-effects models, this allowed for standardized coefficients (e.g., how many standard deviations performance (Drop Error) changes for each standard deviation increase in a predictor variable (e.g. Straightness).

All group comparisons on behavioral metrics (Drop Error, strategies) were performed with nonparametric tests (Kruskall-Wallis to compare all groups, Wilcoxon Rank Sum Test to compare groups pairwise, Wilcoxon signed-rank test to compare values to baseline) as the data was not normally distributed and the assumption of homogeneity of variance was violated.

### Mixed-Effects Models

To leverage the richness of the data, we opted for mixed-effects models (using R lmer in the lme4 package, [Bates et al., 2015] and lme in the nlme package, [Pinheiro et al., 1999]) to estimate how path indices, i.e. strategies, predict performance and neural activity, respectively.

Upon initial inspection of our linear models and results, we realized that relationships were not linear, and thus we opted to use piecewise linear models. First, we estimated breakpoints in the data by running an iterative optimiser (using R optimise), on a function which searched for a change in the direction of the relationship between two variables (predictor and outcome variable) by inserting a breakpoint in the predictor variable at all possible values (excluding the bottom and top 2.5%ile of values to avoid extreme outliers skewing the breakpoint placement), and running the two segments separately (i.e. the predictor was broken down into two sub-variables, before and after the breakpoint). The breakpoint value that lead to the lowest AIC value across all tested models was then used to define a new variable in the data which represented the interaction term of the variable after the breakpoint. Thus for a given predictor variable, we also calculated a new variable using the pmax function, which was zero for all values below the breakpoint, and took on (valuei – breakpoint) for all others (as in Muggeo, 2008), and we refer to this as the ‘interaction term’. If the estimated breakpoint value was at the minimum or maximum of the possible values of the relevant variable, it was deemed invalid, and no breakpoint was established (ie: the relationship between the two variables was estimated as linear). Additionally, we also tested (using ANOVA model comparison) if a linear model outperformed the piecewise model, and only when the piecewise model was significantly better than the linear model, were the breakpoints included in further analysis (see Drop Error breakpoint analysis for example of metric computed, and ANOVA as well as AIC values for model comparison, Supplemental Table 7C).

Next, we defined our mixed-linear models per group, with breakpoints included (if relevant). For example, to understand how performance (outcome) was related to putative strategies, we constructed a model with the following fixed effects: strategy metric and its interaction term. As path metrics are inherently related to each other, where the object is located in the arena and their use changes over the course of the experiment (Supplemental Figure 3 and Figure 3) we establish significance using this reduced model (i.e. each strategy metric separately), and correct for multiple comparisons using FDR. For all models, we report p-values after adjusting for FDR within group. Note that FDR correction is applied to the first leg of the model and the interaction term (which indicates if the breakpoint is significant). Only if the latter is significant after correction did we establish the significance of the second leg. Participants were included as random effects (intercept only), and for the iEEG group, Session was also included as a random effect, as some patients performed multiple sessions of the task, over different days.

For hippocampal activity (and entorhinal/caudate in fMRI samples), we constructed timepoint-by-timepoint models for each group separately. While we acknowledge that brain activity is the driver of behavior, it is only possible to predict one outcome variable as a function of multiple fixed effects, i.e. predictor variables, therefore neural activity was designated the outcome variable. In order to understand not only how neural activity relates to our putative strategy metrics, we wanted to see how it relates as a function of whether the behavior is adaptive (useful), therefore we used the breakpoints for the path indices established for Drop Error, and the associated interaction terms in our model. Thus, our model consisted of path metrics as above with the interaction term. We also ran separate models with the following control variables (given previous reports in the literature): Drop Error (performance), Distance from the Drop Location, Object Distance to Boundary, Speed, Object trial number. We only included those timepoints in this model when the participant was moving (Speed *>*1). Note that the path metrics were either at trial-level (SI,DB,PO), and control predictors were either object-level (Object Distance to Boundary), trial-level (Drop Error, Object trial number) or timepoint-by-timepoint (Distance from the Drop Location, Speed). Due to the high level of autocorrelation in neural time-series data, we used the CAR1 correction package (part of nlme package) in our mixed-level models and confirmed successful autocorrelation removal by inspecting the residuals. All code for path metrics, statistical analysis, and MLMs available on GitHub: XXX.

## Supporting information

Supplementary Results

## Acknowledgements

This project was funded by a grant from the European Research Council (ERC CoG 864164 “Grid Representations”). D. Stawarczyk is Research Associate at the Fonds de la Recherche Scientifique (F.R.S.-FNRS).

