## Supplementary Results for "It’s All In The Journey: Putative Strategies Extracted from Navigation Paths Predict Spatial Memory and Hippocampal Recruitment"

December 14, 2025

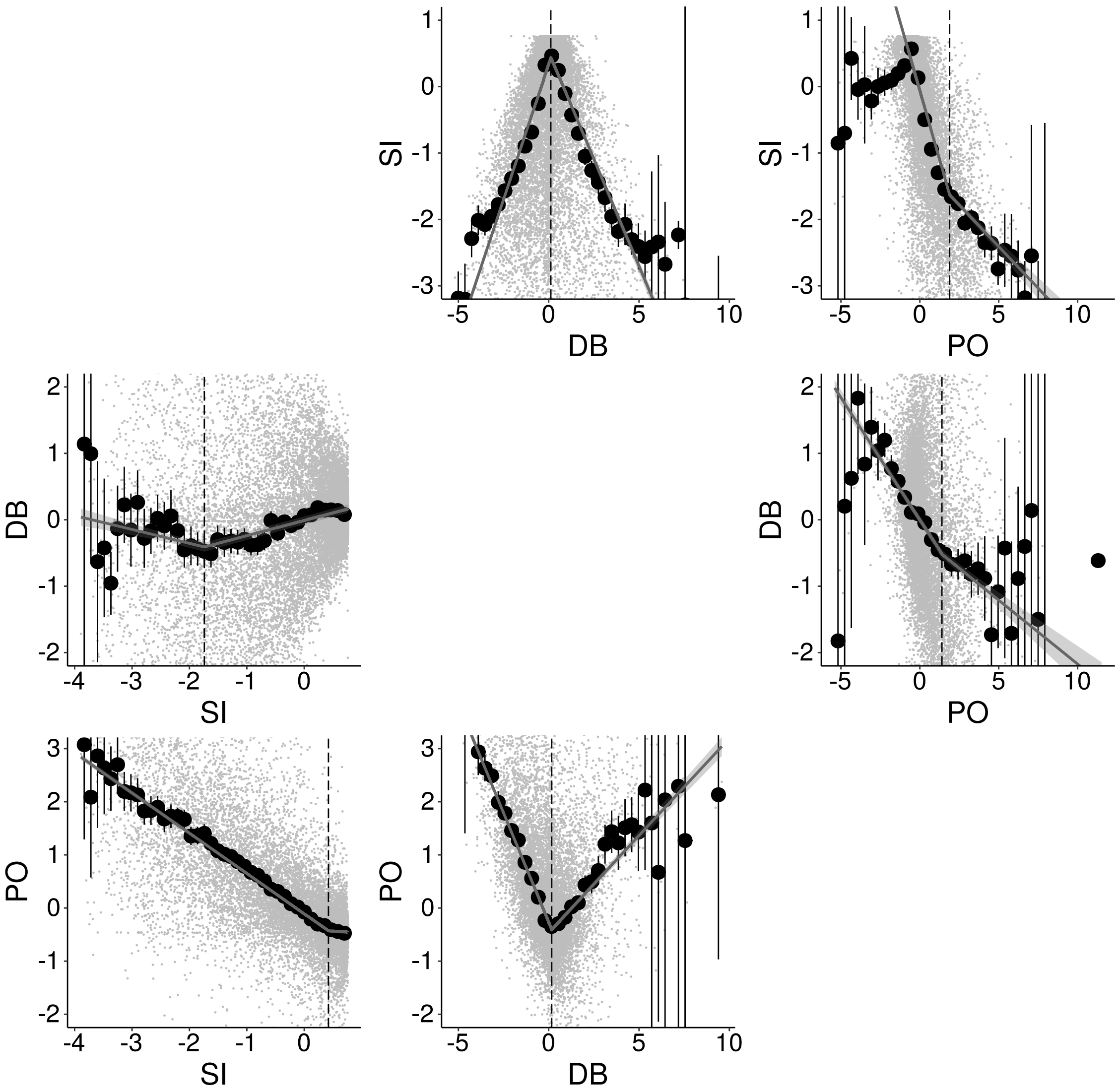


Supplemental Figure 1: Relation between Path Indices (detail for Figure 2B). We used piecewise mixed linear regressions to investigate how our proposed path metrics were related (using data across all participants) and found that all path metrics were significantly related to each other (all B1 = $p<0.001$, with significant breakpoints, and second slope (B2) significant $p<0.001$ in all cases except the slope after the breakpoint in SI $\rightarrow$PO ($p=0.07$), see Supplemental Table 1 for detailed outputs of models). Plots show regression lines with 95% CI overlaid on raw data (grey points), and black dots with confidence intervals are binned data for visualization only.

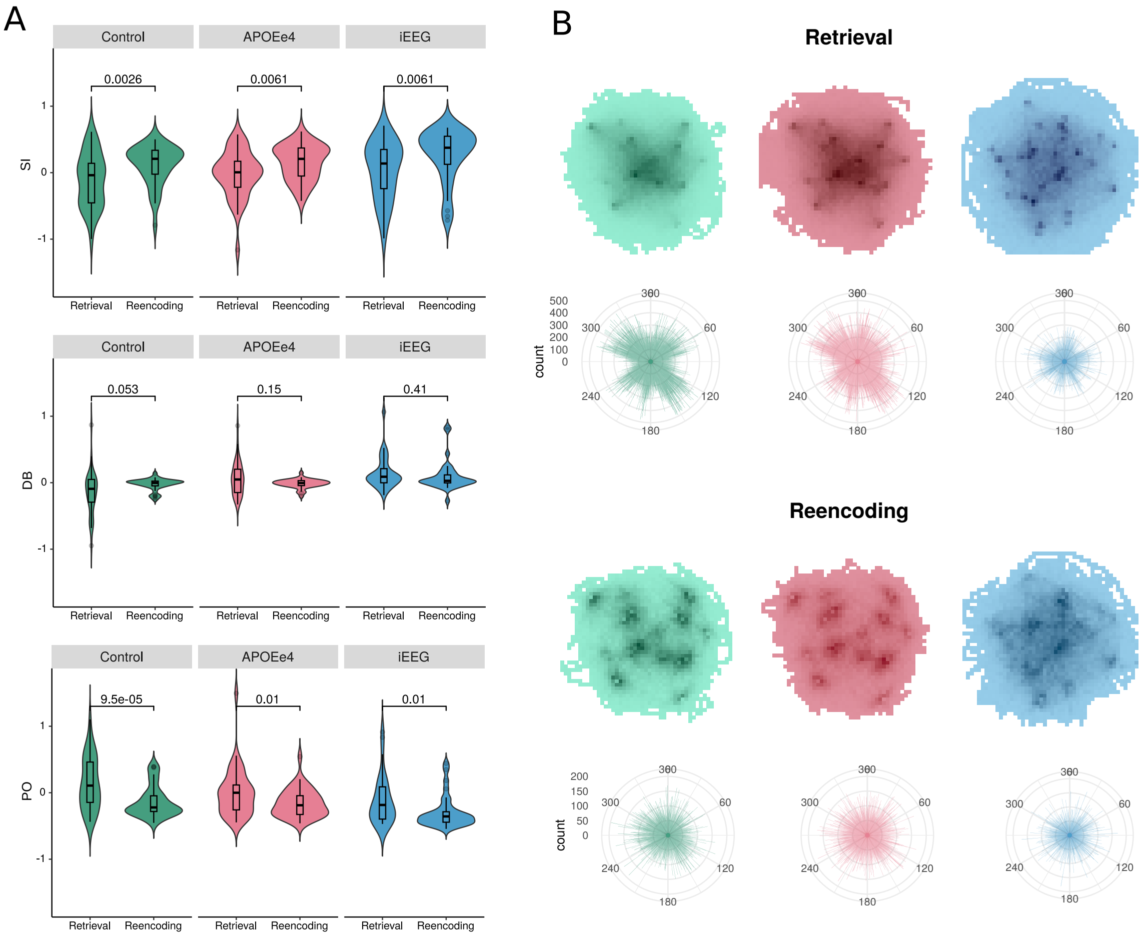


Supplemental Figure 2: Use of Strategies is Unique to Retrieval – Not present during Re-encoding. A) Within group comparison of path indices between retrieval and re-encoding, showing that Straightness (SI) is higher during reencoding (as participants are shown the object and can walk directly towards it), whereas deviating (DB) towards centre is reduced (trend) during reencoding in Controls (note the Control and APOEe4 group show no significant difference from 0 during reencoding, both $p>0.1$, the iEEG group still show significant deviation to the boundary even in reencoding, $p<0.001$). All groups show significantly less overlapping paths during reencoding (PO). [Note z-scoring for re-encoding was based on means/standard deviations from retrieval phase for comparison] B) Heatmap of locations visited and heading directions taken (all timepoints during navigation), plotted by group. During retrieval, map occupancy was more distributed around the arena (note Controls occupy the center more) and heading directions are more clustered in certain directions, which may overlap with distal cues in the arena. During reencoding the locations visited were clustered around the goal locations, and heading was more uniform.

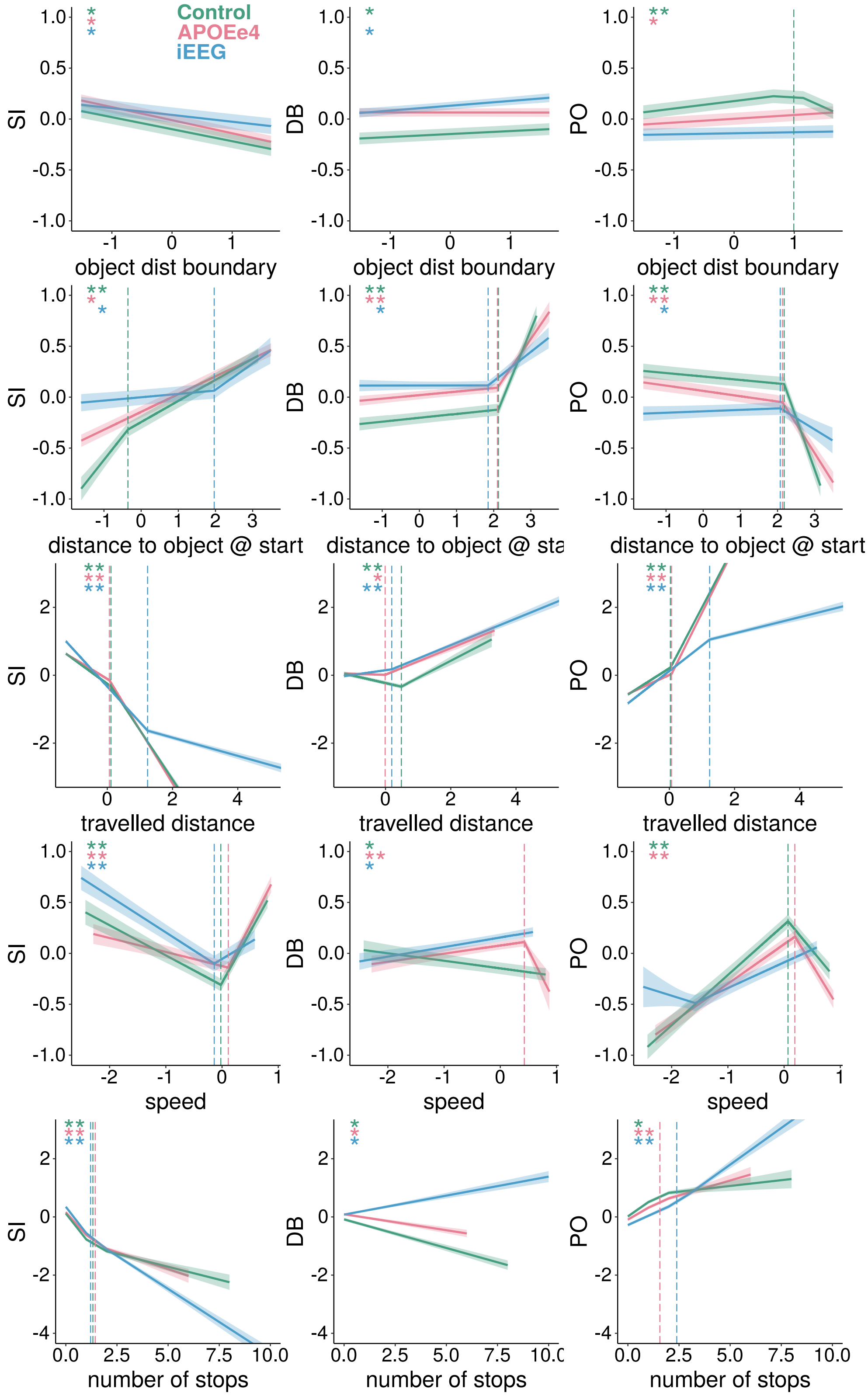

Control
APOEe4
iEEG
SI
DB
PO
object dist boundary
distance to object @ start
distance to object @ sta
travelled distance
speed
number of stops

Supplemental Figure 3: Relation between Path Metrics and Other Behavioral and Experimental Variables. We used piecewise linear regressions to investigate how our proposed path metrics were related to variables including where the object was in the arena (object distance from boundary), how far the goal object was at the start of the trial, the distance travelled in the trial, the speed participants travelled at and the number of times, within a trial, that participants stopped for more than 2 seconds. Overall participants took more straight paths to objects located closer to the boundary and further away from the start point and when they travelled fast. They took less straight paths when they travelled longer distances or stopped many times while navigating. Controls deviated more towards the center when the goal object was near the boundary (perhaps to gauge direction better), and all groups deviated more to the boundary when the goal object was further away from the start location. Increase in distance travelled, speed and number of stops increased deviation – for Controls this meant increased deviation towards the boundary, except for very long travelled distances which were related to increasingly boundary-related deviation. The APOEe4 groups also showed an interesting effect where very high speeds (and increased stopping) were related to deviation towards the center (rather than boundary). Controls and APOEe4s showed increased route matching to object located near the center of the arena, and all groups showed increases with larger travelled distances and number of stops. Interestingly, route matching decreased as the goal object was further from the start location – this may be due to the fact that the overall distance to get to a distant goal would be by definition higher and thus given the distributed nature of the goal locations, the overall percentage of the route that could in practice overlap with all the rest (which may be more bunched up near the center) is just smaller – this also related to the increase seen for objects near the center, which may be an artefact of the increased possibility of a route going through the center. Stars indicate if slope (left star: B1, right star:B2) is significant. Only significant breakpoints are shown. See Supplemental Table 2 for detailed output of models.

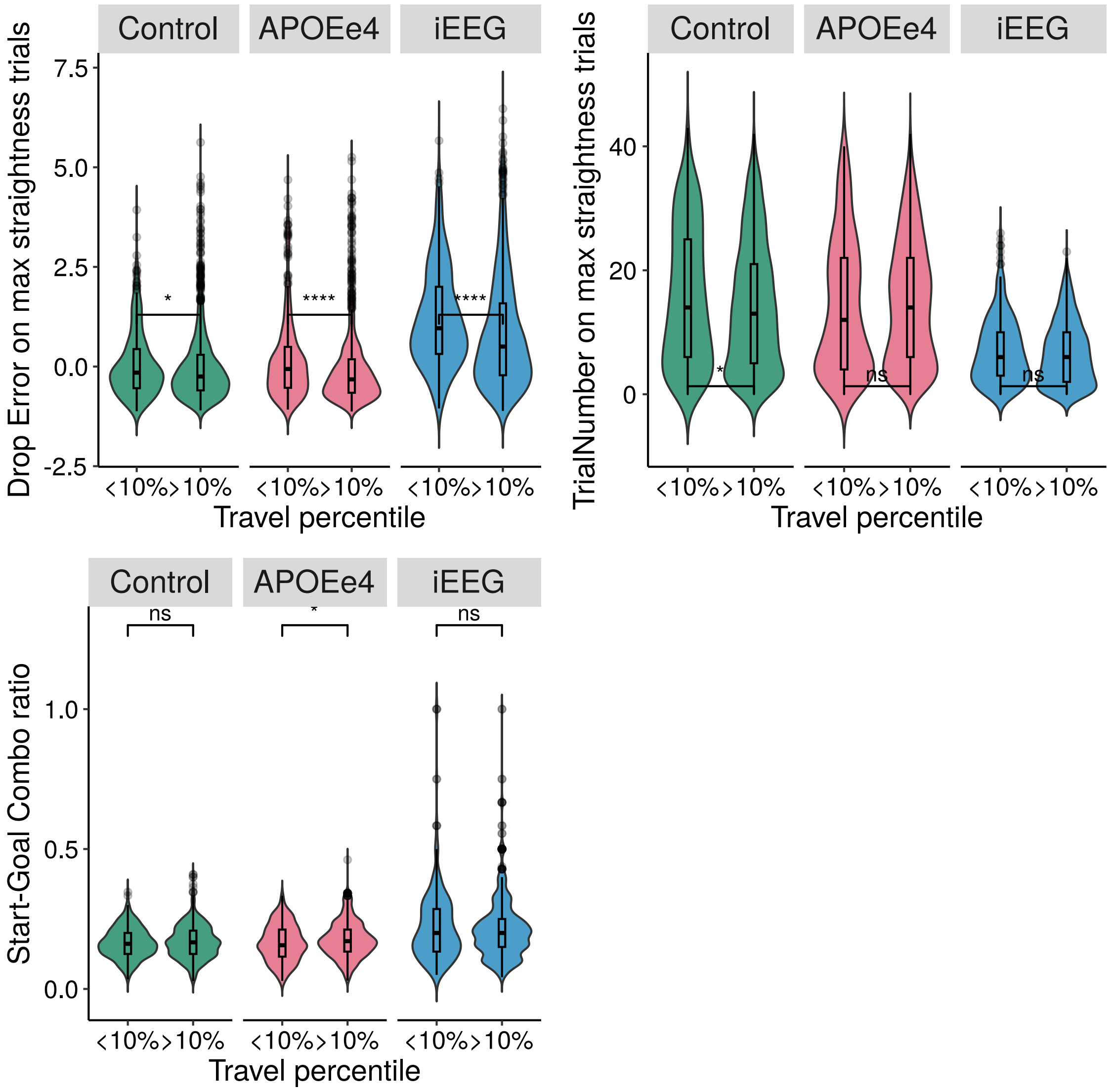


Supplemental Figure 4: Top Left: Very short (<10th percentile of all travelled distance) and simultaneously very straight trials (>90th percentile of all straightness, ie SI>0.9 on raw values) have larger Drop Error compared to longer very straight trials. These are putatively the quick drops participants make when they do not know where the goal object is located and wish to terminate the trial instead of searching. Top Right: Trial number (per object) for max straightness trials for very short and longer paths. It does not seem that these short and very straight trials are confined to earlier or later trials, but occur similarly frequently across the experiment (except in Controls where these short straight trials happen more on later trials). Bottom Left: We calculated the ratio for each object to object pair, ie. The start-goal location combination, in order to understand if the very straight short paths were related to infrequent combinations. This was not clearly the case, although in th APOEe4 group, there was a significant effect of straight-short paths happening on a lower start-goal combo ratio. These analyses do not clearly explain the source of the very short and very straight trials, but performance is reduced on these trials and may be a result of uncertainty.

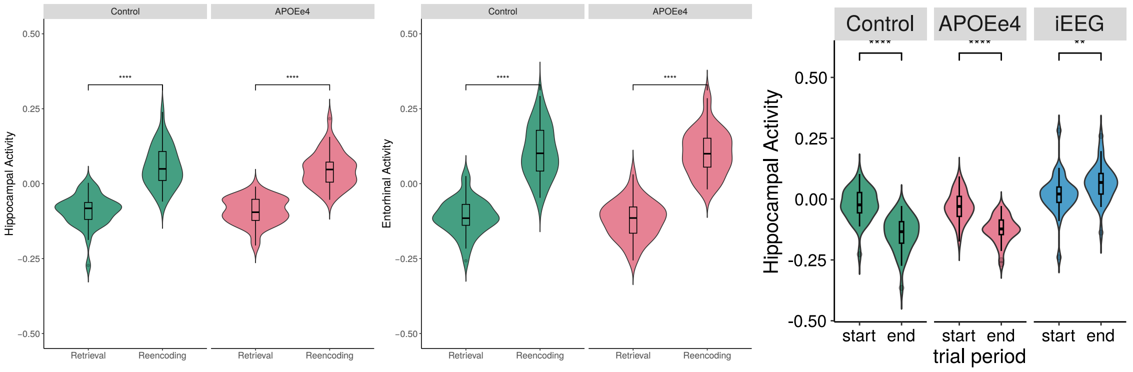


Supplemental Figure 5: Hippocampal and Entorhinal BOLD are both higher during reencoding than retrieval phase. BOLD is also higher at the start of the retrieval trial than at the end, with theta power showing the inverse effect.

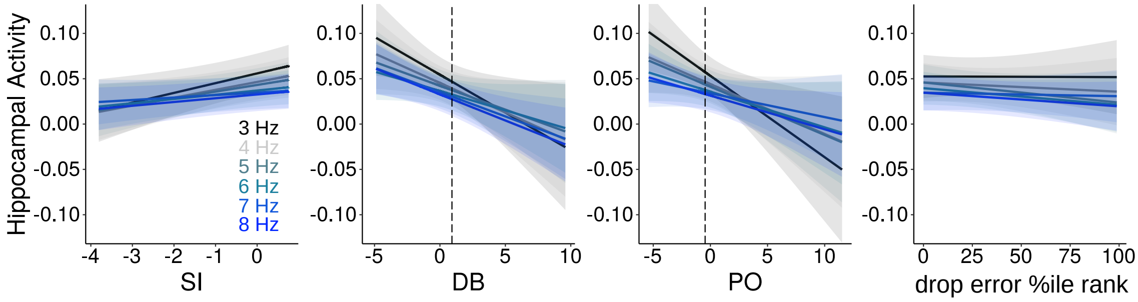


Supplemental Figure 6: Strategy effects on hippocampal theta separated by frequency band. 3Hz was most consistently associated with path metrics. See also Supplemental Table 11 for p-values.

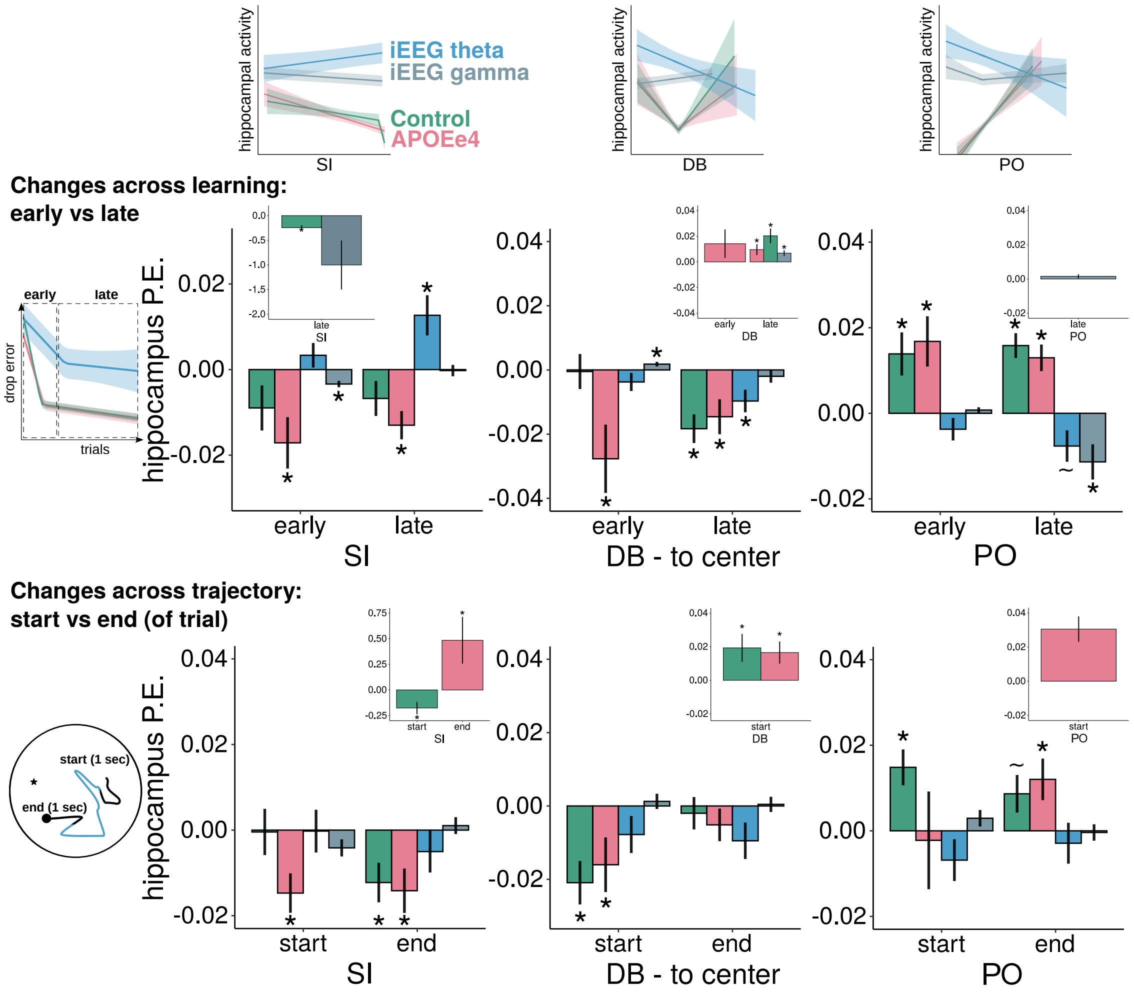


Supplemental Figure 7: Hippocampus and Adaptive Strategy Use by Learning Phase and Within Trials. Schematics of strategy x hippocampal activity results from Figure 6 in the main text as a reminder. Top: inset shows a schematic of the early vs late learning phases in Drop Error. We divided the neural activity into two phases as well to understand the relationship between hippocampal activity and adaptive strategy use over time. Shown are extracted beta coefficients for piecewise regressions run separately for trials in the early and late learning period. Insets show the beta coefficients for the second slope, where applicable. These are only presented if the breakpoint model (i.e. piecewise regression) was significantly better than the linear model. For example, for Straightness, in early learning, a simple linear model explained the relationship between Straightness and hippocampal BOLD activity in Controls. In the late phase, the piecewise model was significantly better, and therefore there is a beta coefficient plotted for the second slope (“B2” in the text). Bottom: inset shows a schematic of the first (”start”) and last (”end”) second of a hypothetical trajectory. Shown are extracted beta coefficients for piecewise regressions run separately for the start and end period of every trial (trials that were not long enough, i.e. less than 1 second for each start/end period or where these segments were overlapping, were not included). Significance (“*”) is shown after correcting for multiple comparisons using FDR. A “ ” indicates that it is a trend effect (after FDR).

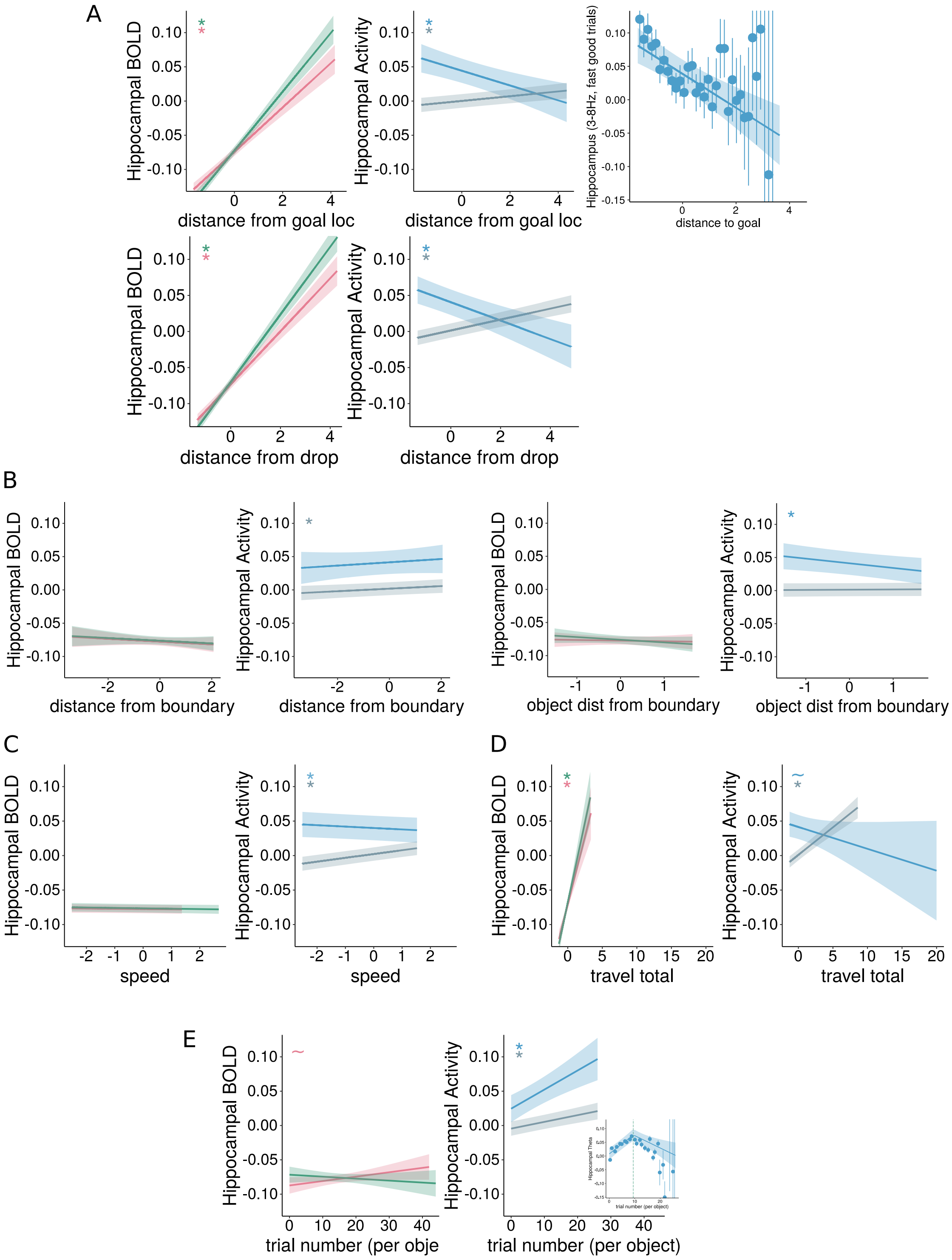

A
Hippocampal BOLD
Hippocampal Activity
distance from goal loc
Hippocampus (3-8Hz, fast good trials)
distance to goal
distance from drop
B
distance from boundary
object dist from boundary
C
speed
D
travel total
E
trial number (per obje
trial number (per object)

Supplemental Figure 8: Neural Activity in Relation to Other Behavioral Variables. We analyzed the effect of several control variables on hippocampal activity during the retrieval period (linear mixed-effects regression only, no interaction terms as with path indices). A) Distance from goal (and drop) location is time-point by time-point measure of the distance participants were from their goal/final drop location (noting that the goal location was hidden and termination of the trial by indicating a drop location was controlled by the participant). Overall, as participants approached their goal/drop, BOLD activity and gamma power decreased and theta power increased. Also, specifically for fast movement with low drop error trials, analyzed as per Liu et al, 2023, we found increased theta power as the final location was approached, which does not align with their findings, but may relate to our hippocampal contacts being predominantly in the anterior hippocampus, Supplemental Figure 10 . B) 'Distance to boundary', is a time-point by time-point measure of the participants' location relative to the boundary, showed decreased gamma power nearer to the boundary but no effects in theta power. 'Object distance from boundary' is the same across the trial as it is a stable value of where the goal object is located in the arena relative to the boundary. We found that theta power was increased for goal locations close to the boundary, similar to previous reports showing that theta power is increased at boundaries - which coincidentally were where the goals were located in those tasks (e.g. Stangl 2020). These results all point to the fact that theta power may be increased at boundaries, but only when the boundaries are relevant for goal location coding. C) 'Speed' was included as time-point by time-point measure of virtual linear speed, and showed a positive relationship with gamma power in the hippocampus, whereas theta power did not change significantly as a function of movement speed in the arena (vs Aghajan/Watrous), but did show a trend in the opposite direction from predicted, i.e. less theta power with increasing speed. D) 'Total travelled distance' which was a measure of the path travelled showed increased gamma power but only a trernd in theta power, although in the opposite direction than previously reported, ie increased theta power dor longer distances (vs Bush), which may be related to the time point at which these effects are measures (cue period/start of the trial vs during movement). E) 'Object Trial Number' represents the number of instances a particular goal object was cued/searched for and can be seen as a measure of learning. Trials to various goal locations were interleaved, so this metric is similar to overall trial number. Hippocampal theta and gamma power increased with repeated visits to goals. When looking at the raw data, we noted that for theta power, this relationship changed sign around the 9th trial, which is around the time that Drop Error showed a breakpoint as a function of 'Object Trial Number' (Figure 1C), therefore as a follow-up, we added the interaction term and found that this piecewise model predicted hippocampal theta better than when 'Object Trial Number' was included as a single linear regressor only. This suggests that theta power increases during early learning only, while gamma power remains elevated throughout navigation. The APOEe4 group showed a trend towards increasing BOLD with learning.

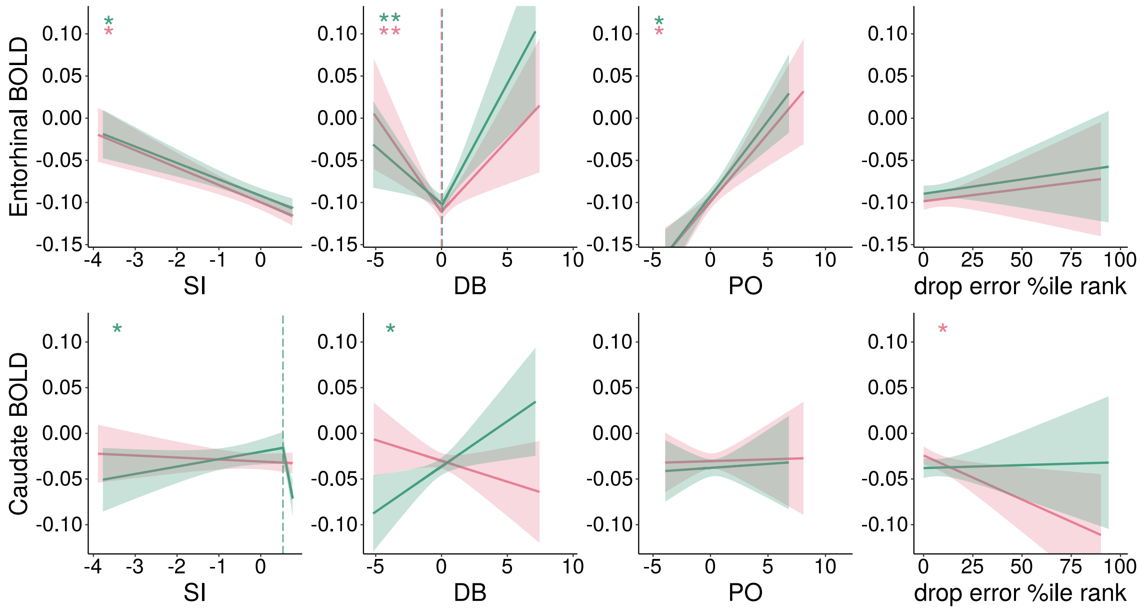


Supplemental Figure 9: Bilateral Entorhinal and Caudate BOLD Activity in Relation to Path Indices and Drop Error. Stars indicate if slope is significant. Only significant breakpoints are shown. See Supplemental Table 12 for detailed output of models.

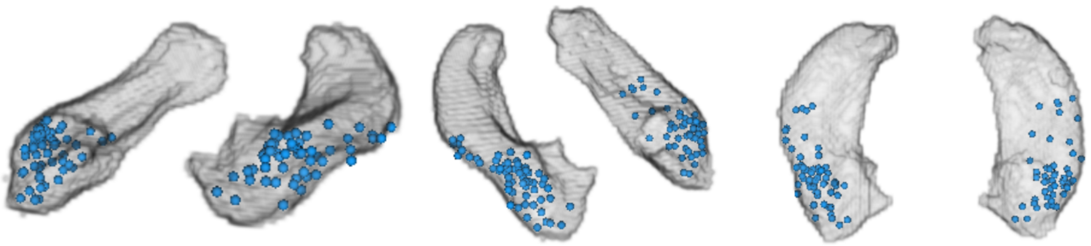

Supplemental Figure 10: Visualization of electrode contact locations (after bipolar referencing) in the hippocampus across all iEEG patients.

Supplementary Table 1: Results of Group Comparisons (Kruskall-Wallis test)

| measure | n | statistic | df | p |
|---|---|---|---|---|
| DropError | 107 | 53.123 | 2 | 0.000 |
| SI | 107 | 2.794 | 2 | 0.247 |
| DB | 107 | 18.16 | 2 | 0.000 |
| PO | 107 | 12.736 | 2 | 0.002 |
| Drop boundary bias | 107 | 15.250 | 2 | 0.000 |

Supplementary Table 2: Results of Follow-up Wilcoxon rank sum tests comparing groups

| measure | group1 | group2 | n1 | n2 | statistic | p | p-adj | signif | effsize |
|---|---|---|---|---|---|---|---|---|---|
| DropError | APOEe4 | Control | 38 | 36 | 655 | 0.759 | 1.000 | ns | 0.036 |
| DropError | APOEe4 | iEEG | 38 | 33 | 71 | 0.000 | 0.000 | **** | 0.761 |
| DropError | Control | iEEG | 36 | 33 | 70 | 0.000 | 0.000 | **** | 0.758 |
| SI | APOEe4 | Control | 38 | 36 | 784 | 0.284 | 0.852 | ns | 0.126 |
| SI | APOEe4 | iEEG | 38 | 33 | 556 | 0.419 | 1.000 | ns | 0.097 |
| SI | Control | iEEG | 36 | 33 | 465 | 0.123 | 0.369 | ns | 0.187 |
| DB | APOEe4 | Control | 38 | 36 | 957 | 0.003 | 0.009 | ** | 0.343 |
| DB | APOEe4 | iEEG | 38 | 33 | 522 | 0.230 | 0.690 | ns | 0.144 |
| DB | Control | iEEG | 36 | 33 | 249 | 0.000 | 0.000 | **** | 0.499 |
| PO | APOEe4 | Control | 38 | 36 | 519 | 0.075 | 0.225 | ns | 0.207 |
| PO | APOEe4 | iEEG | 38 | 33 | 817 | 0.028 | 0.085 | ns | 0.260 |
| PO | Control | iEEG | 36 | 33 | 875 | 0.001 | 0.002 | ** | 0.406 |
| Drop boundary bias | APOEe4 | Control | 38 | 36 | 865 | 0.051 | 0.152 | ns | 0.228 |
| Drop boundary bias | APOEe4 | iEEG | 38 | 33 | 396 | 0.007 | 0.022 | * | 0.316 |
| Drop boundary bias | Control | iEEG | 36 | 33 | 300 | 0.000 | 0.001 | *** | 0.425 |

Supplementary Table 3: Comparison of Path Indices (raw values) to Baseline (0 for DB/PO; 1 for SI; two-sided Wilcoxon signed rank tests) during Retrieval

| group | measure | statistic | p |
|---|---|---|---|
| Control | SI | 0 | 0.000 |
| Control | DB | 185 | 0.019 |
| Control | PO | 666 | 0.000 |
| Control | Drop Boundary bias | 441 | 0.091 |
| APOEe4 | SI | 0 | 0.000 |
| APOEe4 | DB | 489 | 0.087 |
| APOEe4 | PO | 741 | 0.000 |
| APOEe4 | Drop Boundary bias | 640 | 0.000 |
| iEEG | SI | 0 | 0.000 |
| iEEG | DB | 507 | 0.000 |
| iEEG | PO | 516 | 0.000 |
| iEEG | Drop Boundary bias | 516 | 0.000 |

Supplementary Table 4A: Correlation of Path Indices with Age (Spearman's rho)

| group | measure | r(s) | p |
|---|---|---|---|
| Control | Drop Error | 0.36 | 0.0299 |
| APOEe4 | Drop Error | 0.25 | 0.134 |
| iEEG | Drop Error | 0.099 | 0.582 |
| Control | Drop boundary bias | -0.021 | 0.901 |
| APOEe4 | Drop boundary bias | -0.036 | 0.83 |
| iEEG | Drop boundary bias | -0.0097 | 0.957 |
| Control | SI | -0.024 | 0.887 |
| APOEe4 | SI | 0.48 | 0.00235 |
| iEEG | SI | -0.18 | 0.304 |
| Control | DB | -0.17 | 0.336 |
| APOEe4 | DB | -0.2 | 0.235 |
| iEEG | DB | -0.036 | 0.841 |
| Control | PO | 0.006 | 0.972 |
| APOEe4 | PO | -0.49 | 0.00169 |
| iEEG | PO | -0.081 | 0.654 |

Supplementary Table 4B: Gender Effects (Wilcoxon rank-sum test)

| group | measure | gender | gender | N (f) | N(m) | p |
|---|---|---|---|---|---|---|
| Control | Drop Error | f | m | 18 | 18 | 0.126 |
| APOEe4 | Drop Error | f | m | 20 | 18 | 0.264 |
| iEEG | Drop Error | f | m | 20 | 13 | 0.573 |
| Control | Drop boundary bias | f | m | 18 | 18 | 0.839 |
| APOEe4 | Drop boundary bias | f | m | 20 | 18 | 0.806 |
| iEEG | Drop boundary bias | f | m | 20 | 13 | 0.316 |
| Control | SI | f | m | 18 | 18 | 0.355 |
| APOEe4 | SI | f | m | 20 | 18 | 0.828 |
| iEEG | SI | f | m | 20 | 13 | 0.573 |
| Control | DB | f | m | 18 | 18 | 0.542 |
| APOEe4 | DB | f | m | 20 | 18 | 0.196 |
| iEEG | DB | f | m | 20 | 13 | 0.624 |
| Control | PO | f | m | 18 | 18 | 0.339 |
| APOEe4 | PO | f | m | 20 | 18 | 0.654 |
| iEEG | PO | f | m | 20 | 13 | 0.899 |

Supplementary Table 5: Piecewise model outputs of relationship between Path Indices

| DV | IV | b | se | p |
|---|---|---|---|---|
| SI | DB | 0.816 | 0.008 | 0.000 |
| SI | DB interactionbp | -1.462 | 0.013 | 0.000 |
| SI | DB secondslope | -0.646 | 0.009 | 0.000 |
| SI | PO | -0.830 | 0.007 | 0.000 |
| SI | PO interactionbp | 0.579 | 0.019 | 0.000 |
| SI | PO secondslope | -0.251 | 0.015 | 0.000 |
| DB | PO | -0.368 | 0.011 | 0.000 |
| DB | PO interactionbp | 0.176 | 0.024 | 0.000 |
| DB | PO secondslope | -0.192 | 0.017 | 0.000 |
| DB | SI | -0.208 | 0.033 | 0.000 |
| DB | SI interactionbp | 0.436 | 0.038 | 0.000 |
| DB | SI secondslope | 0.228 | 0.010 | 0.000 |
| PO | SI | -0.757 | 0.007 | 0.000 |
| PO | SI interactionbp | 0.681 | 0.047 | 0.000 |
| PO | SI secondslope | -0.077 | 0.043 | 0.076 |
| PO | DB | -0.824 | 0.008 | 0.000 |
| PO | DB interactionbp | 1.190 | 0.014 | 0.000 |
| PO | DB secondslope | 0.366 | 0.010 | 0.000 |

Supplementary Table 6: Piecewise model outputs of relationship between Path Indices and other Behavioural and Experimental Variables. P-values were corrected using FDR, which are the values reported in the text and figures. The significance of the second slope was only established if the breakpoint itself was significant after correction, therefore these values are not corrected for FDR. As an example, in the iEEG group, PO was predicted by distance to the goal location from the start of the trial ("Dist to Object trlstart"). The left slope of the breakpoint was not significant (p fdr=0.529), the breakpoint was significant (p fdr=0.02), and the right slope was significant (p = 0.011).

| group | DV | IV | b | se | p | p FDR |
|---|---|---|---|---|---|---|
| Control | SI | Object TrialNum | -0.128 | 0.025 | 0.000 | 0.000 |
| Control | SI | Object TrialNum interactionbp | 0.130 | 0.025 | 0.000 | 0.000 |
| Control | SI | Object TrialNum secondslope | 0.002 | 0.001 | 0.176 | 0.176 |
| Control | SI | Dist Obj From Boundary | -0.118 | 0.010 | 0.000 | 0.000 |
| Control | SI | Dist to Object trlstart | 0.461 | 0.081 | 0.000 | 0.000 |
| Control | SI | Dist to Object trlstart interactionbp | -0.254 | 0.087 | 0.003 | 0.004 |
| Control | SI | Dist to Object trlstart secondslope | 0.207 | 0.012 | 0.000 | 0.000 |
| Control | SI | Total Traveled Dist | -0.694 | 0.028 | 0.000 | 0.000 |
| Control | SI | Total Traveled Dist interactionbp | -0.758 | 0.063 | 0.000 | 0.000 |
| Control | SI | Total Traveled Dist secondslope | -1.452 | 0.047 | 0.000 | 0.000 |
| Control | SI | Speed | -0.295 | 0.049 | 0.000 | 0.000 |
| Control | SI | Speed interactionbp | 1.295 | 0.099 | 0.000 | 0.000 |
| Control | SI | Speed secondslope | 1.000 | 0.068 | 0.000 | 0.000 |
| Control | SI | N stopped | -0.891 | 0.027 | 0.000 | 0.000 |
| Control | SI | N stopped interactionbp | 0.714 | 0.057 | 0.000 | 0.000 |
| Control | SI | N stopped secondslope | -0.177 | 0.040 | 0.000 | 0.000 |
| Control | DB | Object TrialNum | -0.179 | 0.043 | 0.000 | 0.000 |
| Control | DB | Object TrialNum interactionbp | 0.184 | 0.043 | 0.000 | 0.000 |
| Control | DB | Object TrialNum secondslope | 0.005 | 0.001 | 0.000 | 0.000 |
| Control | DB | Dist Obj From Boundary | 0.029 | 0.011 | 0.006 | 0.007 |
| Control | DB | Dist to Object trlstart | 0.038 | 0.013 | 0.004 | 0.004 |
| Control | DB | Dist to Object trlstart interactionbp | 0.862 | 0.090 | 0.000 | 0.000 |
| Control | DB | Dist to Object trlstart secondslope | 0.899 | 0.084 | 0.000 | 0.000 |
| Control | DB | Total Traveled Dist | -0.215 | 0.027 | 0.000 | 0.000 |
| Control | DB | Total Traveled Dist interactionbp | 0.716 | 0.091 | 0.000 | 0.000 |
| Control | DB | Total Traveled Dist secondslope | 0.500 | 0.079 | 0.000 | 0.000 |
| Control | DB | Speed | -0.073 | 0.032 | 0.021 | 0.021 |
| Control | DB | N stopped | -0.197 | 0.020 | 0.000 | 0.000 |
| Control | PO | Object TrialNum | 0.024 | 0.004 | 0.000 | 0.000 |
| Control | PO | Object TrialNum interactionbp | -0.028 | 0.005 | 0.000 | 0.000 |
| Control | PO | Object TrialNum secondslope | -0.004 | 0.002 | 0.030 | 0.030 |
| Control | PO | Dist Obj From Boundary | 0.073 | 0.015 | 0.000 | 0.000 |
| Control | PO | Dist Obj From Boundary interactionbp | -0.338 | 0.069 | 0.000 | 0.000 |
| Control | PO | Dist Obj From Boundary secondslope | -0.265 | 0.060 | 0.000 | 0.000 |
| Control | PO | Dist to Object trlstart | -0.034 | 0.013 | 0.008 | 0.009 |
| Control | PO | Dist to Object trlstart interactionbp | -0.990 | 0.098 | 0.000 | 0.000 |
| Control | PO | Dist to Object trlstart secondslope | -1.024 | 0.092 | 0.000 | 0.000 |
| Control | PO | Total Traveled Dist | 0.609 | 0.029 | 0.000 | 0.000 |
| Control | PO | Total Traveled Dist interactionbp | 1.223 | 0.061 | 0.000 | 0.000 |
| Control | PO | Total Traveled Dist secondslope | 1.832 | 0.043 | 0.000 | 0.000 |
| Control | PO | Speed | 0.493 | 0.046 | 0.000 | 0.000 |
| Control | PO | Speed interactionbp | -1.160 | 0.113 | 0.000 | 0.000 |
| Control | PO | Speed secondslope | -0.666 | 0.086 | 0.000 | 0.000 |
| Control | PO | N stopped | 0.492 | 0.028 | 0.000 | 0.000 |
| Control | PO | N stopped interactionbp | -0.412 | 0.067 | 0.000 | 0.000 |
| Control | PO | N stopped secondslope | 0.079 | 0.051 | 0.120 | 0.120 |
| APOEe4 | SI | Object TrialNum | -0.005 | 0.001 | 0.000 | 0.000 |
| APOEe4 | SI | Dist Obj From Boundary | -0.129 | 0.010 | 0.000 | 0.000 |
| APOEe4 | SI | Dist to Object trlstart | 0.175 | 0.010 | 0.000 | 0.000 |

| Continuation of Table S6 | | | | | | |
|---|---|---|---|---|---|---|
| group | DV | IV | b | se | p | p FDR |
| APOEe4 | SI | Total Traveled Dist | -0.581 | 0.026 | 0.000 | 0.000 |
| APOEe4 | SI | Total Traveled Dist interactionbp | -0.987 | 0.056 | 0.000 | 0.000 |
| APOEe4 | SI | Total Traveled Dist secondslope | -1.568 | 0.041 | 0.000 | 0.000 |
| APOEe4 | SI | Speed | -0.137 | 0.038 | 0.000 | 0.000 |
| APOEe4 | SI | Speed interactionbp | 1.207 | 0.109 | 0.000 | 0.000 |
| APOEe4 | SI | Speed secondslope | 1.070 | 0.089 | 0.000 | 0.000 |
| APOEe4 | SI | N stopped | -0.782 | 0.025 | 0.000 | 0.000 |
| APOEe4 | SI | N stopped interactionbp | 0.549 | 0.068 | 0.000 | 0.000 |
| APOEe4 | SI | N stopped secondslope | -0.233 | 0.054 | 0.000 | 0.000 |
| APOEe4 | DB | Object TrialNum | 0.007 | 0.001 | 0.000 | 0.000 |
| APOEe4 | DB | Object TrialNum interactionbp | -0.042 | 0.011 | 0.000 | 0.000 |
| APOEe4 | DB | Object TrialNum secondslope | -0.035 | 0.011 | 0.001 | 0.001 |
| APOEe4 | DB | Dist Obj From Boundary | -0.001 | 0.010 | 0.958 | 0.958 |
| APOEe4 | DB | Dist to Object trlstart | 0.034 | 0.013 | 0.007 | 0.008 |
| APOEe4 | DB | Dist to Object trlstart interactionbp | 0.501 | 0.077 | 0.000 | 0.000 |
| APOEe4 | DB | Dist to Object trlstart secondslope | 0.535 | 0.071 | 0.000 | 0.000 |
| APOEe4 | DB | Total Traveled Dist | -0.031 | 0.034 | 0.365 | 0.378 |
| APOEe4 | DB | Total Traveled Dist interactionbp | 0.420 | 0.068 | 0.000 | 0.000 |
| APOEe4 | DB | Total Traveled Dist secondslope | 0.389 | 0.047 | 0.000 | 0.000 |
| APOEe4 | DB | Speed | 0.080 | 0.033 | 0.016 | 0.017 |
| APOEe4 | DB | Speed interactionbp | -1.173 | 0.434 | 0.007 | 0.008 |
| APOEe4 | DB | Speed secondslope | -1.093 | 0.423 | 0.010 | 0.010 |
| APOEe4 | DB | N stopped | -0.110 | 0.021 | 0.000 | 0.000 |
| APOEe4 | PO | Object TrialNum | 0.015 | 0.003 | 0.000 | 0.000 |
| APOEe4 | PO | Object TrialNum interactionbp | -0.015 | 0.004 | 0.001 | 0.001 |
| APOEe4 | PO | Object TrialNum secondslope | 0.000 | 0.002 | 0.855 | 0.855 |
| APOEe4 | PO | Dist Obj From Boundary | 0.037 | 0.009 | 0.000 | 0.000 |
| APOEe4 | PO | Dist to Object trlstart | -0.051 | 0.011 | 0.000 | 0.000 |
| APOEe4 | PO | Dist to Object trlstart interactionbp | -0.527 | 0.073 | 0.000 | 0.000 |
| APOEe4 | PO | Dist to Object trlstart secondslope | -0.578 | 0.068 | 0.000 | 0.000 |
| APOEe4 | PO | Total Traveled Dist | 0.440 | 0.024 | 0.000 | 0.000 |
| APOEe4 | PO | Total Traveled Dist interactionbp | 1.491 | 0.052 | 0.000 | 0.000 |
| APOEe4 | PO | Total Traveled Dist secondslope | 1.931 | 0.038 | 0.000 | 0.000 |
| APOEe4 | PO | Speed | 0.387 | 0.035 | 0.000 | 0.000 |
| APOEe4 | PO | Speed interactionbp | -1.290 | 0.126 | 0.000 | 0.000 |
| APOEe4 | PO | Speed secondslope | -0.902 | 0.109 | 0.000 | 0.000 |
| APOEe4 | PO | N stopped | 0.410 | 0.025 | 0.000 | 0.000 |
| APOEe4 | PO | N stopped interactionbp | -0.205 | 0.075 | 0.006 | 0.007 |
| APOEe4 | PO | N stopped secondslope | 0.205 | 0.061 | 0.001 | 0.001 |
| iEEG | SI | Object TrialNum | 0.004 | 0.003 | 0.247 | 0.314 |
| iEEG | SI | Dist Obj From Boundary | -0.067 | 0.015 | 0.000 | 0.000 |
| iEEG | SI | Dist to Object trlstart | 0.032 | 0.019 | 0.088 | 0.120 |
| iEEG | SI | Dist to Object trlstart interactionbp | 0.228 | 0.087 | 0.008 | 0.013 |
| iEEG | SI | Dist to Object trlstart secondslope | 0.260 | 0.077 | 0.001 | 0.001 |
| iEEG | SI | Total Traveled Dist | -1.052 | 0.024 | 0.000 | 0.000 |
| iEEG | SI | Total Traveled Dist interactionbp | 0.783 | 0.043 | 0.000 | 0.000 |
| iEEG | SI | Total Traveled Dist secondslope | -0.269 | 0.030 | 0.000 | 0.000 |
| iEEG | SI | Speed | -0.355 | 0.047 | 0.000 | 0.000 |
| iEEG | SI | Speed interactionbp | 0.683 | 0.126 | 0.000 | 0.000 |
| iEEG | SI | Speed secondslope | 0.328 | 0.099 | 0.001 | 0.001 |
| iEEG | SI | N stopped | -0.907 | 0.037 | 0.000 | 0.000 |
| iEEG | SI | N stopped interactionbp | 0.452 | 0.056 | 0.000 | 0.000 |
| iEEG | SI | N stopped secondslope | -0.455 | 0.030 | 0.000 | 0.000 |
| iEEG | DB | Object TrialNum | 0.001 | 0.003 | 0.812 | 0.874 |
| iEEG | DB | Dist Obj From Boundary | 0.047 | 0.014 | 0.001 | 0.001 |
| iEEG | DB | Dist to Object trlstart | 0.000 | 0.019 | 0.983 | 0.983 |
| iEEG | DB | Dist to Object trlstart interactionbp | 0.286 | 0.075 | 0.000 | 0.000 |
| iEEG | DB | Dist to Object trlstart secondslope | 0.287 | 0.066 | 0.000 | 0.000 |
| iEEG | DB | Total Traveled Dist | 0.137 | 0.036 | 0.000 | 0.000 |

| Continuation of Table S6 | | | | | | |
|---|---|---|---|---|---|---|
| group | DV | IV | b | se | p | p FDR |
| iEEG | DB | Total Traveled Dist interactionbp | 0.256 | 0.050 | 0.000 | 0.000 |
| iEEG | DB | Total Traveled Dist secondslope | 0.393 | 0.027 | 0.000 | 0.000 |
| iEEG | DB | Speed | 0.093 | 0.032 | 0.003 | 0.006 |
| iEEG | DB | N stopped | 0.131 | 0.019 | 0.000 | 0.000 |
| iEEG | PO | Object TrialNum | 0.000 | 0.003 | 0.906 | 0.939 |
| iEEG | PO | Dist Obj From Boundary | 0.010 | 0.015 | 0.485 | 0.543 |
| iEEG | PO | Dist to Object trlstart | 0.014 | 0.019 | 0.454 | 0.529 |
| iEEG | PO | Dist to Object trlstart interactionbp | -0.238 | 0.096 | 0.014 | 0.020 |
| iEEG | PO | Dist to Object trlstart secondslope | -0.224 | 0.088 | 0.011 | 0.011 |
| iEEG | PO | Total Traveled Dist | 0.750 | 0.027 | 0.000 | 0.000 |
| iEEG | PO | Total Traveled Dist interactionbp | -0.511 | 0.048 | 0.000 | 0.000 |
| iEEG | PO | Total Traveled Dist secondslope | 0.239 | 0.034 | 0.000 | 0.000 |
| iEEG | PO | Speed | -0.173 | 0.230 | 0.453 | 0.529 |
| iEEG | PO | Speed interactionbp | 0.424 | 0.250 | 0.090 | 0.120 |
| iEEG | PO | N stopped | 0.318 | 0.029 | 0.000 | 0.000 |
| iEEG | PO | N stopped interactionbp | 0.186 | 0.066 | 0.005 | 0.008 |
| iEEG | PO | N stopped secondslope | 0.504 | 0.050 | 0.000 | 0.000 |

Supplementary Table 7: Piecewise model outputs for Drop Error. P-values were corrected using FDR, which are the values reported in the text and figures. The significance of the second slope was only established if the breakpoint itself was significant after correction, therefore these values are not corrected for FDR.

| group | DV | IV | b | se | p | p FDR |
|---|---|---|---|---|---|---|
| Control | Drop Error | SI | -0.596 | 0.159 | 0.000 | 0.000 |
| Control | Drop Error | SI interactionbp | 15.536 | 1.807 | 0.000 | 0.000 |
| Control | Drop Error | SI secondslope | 14.940 | 1.719 | 0.000 | 0.000 |
| Control | Drop Error | DB | 0.745 | 0.177 | 0.000 | 0.000 |
| Control | Drop Error | DB interactionbp | -0.942 | 0.335 | 0.005 | 0.005 |
| Control | Drop Error | DB secondslope | -0.197 | 0.245 | 0.422 | 0.422 |
| Control | Drop Error | PO | -1.451 | 0.273 | 0.000 | 0.000 |
| Control | Drop Error | PO interactionbp | 1.388 | 0.443 | 0.002 | 0.002 |
| Control | Drop Error | PO secondslope | -0.063 | 0.236 | 0.791 | 0.791 |
| Control | Drop Error | ObjectTrialNum | -3.931 | 0.107 | 0.000 | 0.000 |
| Control | Drop Error | ObjectTrialNum interactionbp | 3.819 | 0.115 | 0.000 | 0.000 |
| Control | Drop Error | ObjectTrialNum secondslope | -0.112 | 0.015 | 0.000 | 0.000 |
| APOEe4 | Drop Error | SI | -0.376 | 0.140 | 0.007 | 0.008 |
| APOEe4 | Drop Error | SI interactionbp | 68.169 | 5.489 | 0.000 | 0.000 |
| APOEe4 | Drop Error | SI secondslope | 67.793 | 5.435 | 0.000 | 0.000 |
| APOEe4 | Drop Error | DB | 0.419 | 0.208 | 0.044 | 0.044 |
| APOEe4 | Drop Error | DB interactionbp | -1.091 | 0.310 | 0.000 | 0.001 |
| APOEe4 | Drop Error | DB secondslope | -0.671 | 0.184 | 0.000 | 0.000 |
| APOEe4 | Drop Error | PO | -1.751 | 0.330 | 0.000 | 0.000 |
| APOEe4 | Drop Error | PO interactionbp | 1.848 | 0.463 | 0.000 | 0.000 |
| APOEe4 | Drop Error | PO secondslope | 0.097 | 0.209 | 0.641 | 0.641 |
| APOEe4 | Drop Error | ObjectTrialNum | -3.179 | 0.099 | 0.000 | 0.000 |
| APOEe4 | Drop Error | ObjectTrialNum interactionbp | 3.047 | 0.107 | 0.000 | 0.000 |
| APOEe4 | Drop Error | ObjectTrialNum secondslope | -0.132 | 0.015 | 0.000 | 0.000 |
| iEEG | Drop Error | SI | -0.877 | 0.321 | 0.006 | 0.013 |
| iEEG | Drop Error | SI interactionbp | 428.199 | 80.942 | 0.000 | 0.000 |
| iEEG | Drop Error | SI secondslope | 427.322 | 80.837 | 0.000 | 0.000 |
| iEEG | Drop Error | DB | -1.177 | 0.464 | 0.011 | 0.015 |
| iEEG | Drop Error | DB interactionbp | 2.137 | 0.900 | 0.018 | 0.018 |
| iEEG | Drop Error | DB secondslope | 0.960 | 0.644 | 0.136 | 0.136 |
| iEEG | Drop Error | PO | 2.429 | 0.927 | 0.009 | 0.014 |
| iEEG | Drop Error | PO interactionbp | -2.455 | 1.038 | 0.018 | 0.018 |
| iEEG | Drop Error | PO secondslope | -0.026 | 0.343 | 0.940 | 0.940 |
| iEEG | Drop Error | ObjectTrialNum | -1.191 | 0.106 | 0.000 | 0.000 |
| iEEG | Drop Error | ObjectTrialNum interactionbp | 1.061 | 0.215 | 0.000 | 0.000 |
| iEEG | Drop Error | ObjectTrialNum secondslope | -0.129 | 0.143 | 0.365 | 0.365 |

Supplementary Table 7B: Piecewise model outputs for Drop Error with other path metrics as covariates, without interaction terms. No FDR correction applied.

| group | DV | IV | b | se | p |
|---|---|---|---|---|---|
| Control | DropError | SI | -1.611 | 0.214 | 0.000 |
| Control | DropError | SI interactionbp | 16.538 | 1.807 | 0.000 |
| Control | DropError | DB | 0.072 | 0.140 | 0.605 |
| Control | DropError | PO | -1.251 | 0.194 | 0.000 |
| APOE | DropError | SI | -1.067 | 0.199 | 0.000 |
| APOE | DropError | SI interactionbp | 70.226 | 5.504 | 0.000 |
| APOE | DropError | DB | -0.310 | 0.124 | 0.012 |
| APOE | DropError | PO | -0.989 | 0.197 | 0.000 |
| iEEG | DropError | SI | -0.847 | 0.387 | 0.029 |
| iEEG | DropError | SI interactionbp | 425.403 | 81.147 | 0.000 |
| iEEG | DropError | DB | -0.334 | 0.321 | 0.299 |
| iEEG | DropError | PO | 0.058 | 0.371 | 0.876 |
| Control | DropError | DB | 0.453 | 0.235 | 0.053 |
| Control | DropError | DB interactionbp | -0.815 | 0.411 | 0.047 |
| Control | DropError | SI | -0.800 | 0.201 | 0.000 |
| Control | DropError | PO | -1.050 | 0.198 | 0.000 |
| APOE | DropError | DB | 0.085 | 0.263 | 0.747 |
| APOE | DropError | DB interactionbp | -0.709 | 0.397 | 0.074 |
| APOE | DropError | SI | -0.339 | 0.204 | 0.098 |
| APOE | DropError | PO | -0.719 | 0.201 | 0.000 |
| iEEG | DropError | DB | -1.163 | 0.505 | 0.021 |
| iEEG | DropError | DB interactionbp | 2.159 | 1.033 | 0.037 |
| iEEG | DropError | SI | 0.107 | 0.401 | 0.790 |
| iEEG | DropError | PO | 0.140 | 0.371 | 0.706 |
| Control | DropError | PO | -2.286 | 0.347 | 0.000 |
| Control | DropError | PO interactionbp | 1.855 | 0.456 | 0.000 |
| Control | DropError | SI | -0.850 | 0.194 | 0.000 |
| Control | DropError | DB | 0.040 | 0.141 | 0.778 |
| APOE | DropError | PO | -2.054 | 0.366 | 0.000 |
| APOE | DropError | PO interactionbp | 1.923 | 0.464 | 0.000 |
| APOE | DropError | SI | -0.219 | 0.189 | 0.246 |
| APOE | DropError | DB | -0.355 | 0.125 | 0.004 |
| iEEG | DropError | PO | 2.490 | 0.936 | 0.008 |
| iEEG | DropError | PO interactionbp | -2.959 | 1.104 | 0.007 |
| iEEG | DropError | SI | -0.576 | 0.392 | 0.141 |
| iEEG | DropError | DB | -0.344 | 0.322 | 0.286 |

Supplementary Table 7C: Established breakpoints and model comparisons for linear vs breakpoint models for Drop Error .

| group | PI | breakpoint | p linear | p anova linear vs breakpoint | AIC difference |
|---|---|---|---|---|---|
| Control | SI | 0.532 | 0.116 | 0.000 | -71.404 |
| Control | DB | 0.055 | 0.002 | 0.005 | -5.909 |
| Control | PO | 0.740 | 0.000 | 0.002 | -7.843 |
| Control | Object TrialNum | 5.694 | 0.000 | 0.000 | -1041.949 |
| APOEe4 | SI | 0.688 | 0.013 | 0.000 | -150.928 |
| APOEe4 | DB | -0.020 | 0.136 | 0.000 | -10.399 |
| APOEe4 | PO | 0.270 | 0.000 | 0.000 | -13.886 |
| APOEe4 | Object TrialNum | 5.857 | 0.000 | 0.000 | -776.603 |
| iEEG | SI | 0.747 | 0.292 | 0.000 | -25.903 |
| iEEG | DB | 0.918 | 0.240 | 0.018 | -3.630 |
| iEEG | PO | -0.460 | 0.239 | 0.018 | -3.596 |
| iEEG | Object TrialNum | 9.470 | 0.000 | 0.000 | -22.411 |

Supplementary Table 8: Piecewise model outputs for Drop Error – early vs late learning.

| group | learning phase | IV | b | se | p | p FDR |
|---|---|---|---|---|---|---|
| Control | early | SI | -0.633 | 0.557 | 0.256 | 0.256 |
| Control | early | SI interactionbp | 22.342 | 6.360 | 0.000 | 0.001 |
| Control | early | SI secondslope | 21.710 | 6.026 | 0.000 | 0.000 |
| Control | early | DB | 0.723 | 0.468 | 0.122 | 0.138 |
| Control | early | PO | -1.211 | 0.448 | 0.007 | 0.009 |
| Control | later | SI | -0.370 | 0.112 | 0.001 | 0.002 |
| Control | later | SI interactionbp | 5.751 | 1.271 | 0.000 | 0.000 |
| Control | later | SI secondslope | 5.381 | 1.211 | 0.000 | 0.000 |
| Control | later | DB | 0.246 | 0.087 | 0.005 | 0.009 |
| Control | later | PO | -0.733 | 0.189 | 0.000 | 0.000 |
| Control | later | PO interactionbp | 0.823 | 0.304 | 0.007 | 0.009 |
| Control | later | PO secondslope | 0.090 | 0.163 | 0.582 | 0.582 |
| APOEe4 | early | SI | -1.592 | 0.465 | 0.001 | 0.001 |
| APOEe4 | early | SI interactionbp | 79.097 | 17.030 | 0.000 | 0.000 |
| APOEe4 | early | SI secondslope | 77.506 | 16.826 | 0.000 | 0.000 |
| APOEe4 | early | DB | -1.094 | 0.469 | 0.020 | 0.025 |
| APOEe4 | early | PO | 0.434 | 0.452 | 0.337 | 0.375 |
| APOEe4 | later | SI | -0.076 | 0.110 | 0.487 | 0.487 |
| APOEe4 | later | SI interactionbp | 35.653 | 4.424 | 0.000 | 0.000 |
| APOEe4 | later | SI secondslope | 35.576 | 4.383 | 0.000 | 0.000 |
| APOEe4 | later | DB | 0.567 | 0.160 | 0.000 | 0.001 |
| APOEe4 | later | DB interactionbp | -0.729 | 0.235 | 0.002 | 0.003 |
| APOEe4 | later | DB secondslope | -0.162 | 0.137 | 0.235 | 0.235 |
| APOEe4 | later | PO | -1.421 | 0.253 | 0.000 | 0.000 |
| APOEe4 | later | PO interactionbp | 1.381 | 0.357 | 0.000 | 0.000 |
| APOEe4 | later | PO secondslope | -0.040 | 0.163 | 0.808 | 0.808 |
| iEEG | early | SI | -0.725 | 0.393 | 0.065 | 0.114 |
| iEEG | early | SI interactionbp | 330.091 | 99.348 | 0.001 | 0.004 |
| iEEG | early | SI secondslope | 329.366 | 99.213 | 0.001 | 0.001 |
| iEEG | early | DB | -0.709 | 0.400 | 0.076 | 0.114 |
| iEEG | early | PO | 0.626 | 0.370 | 0.091 | 0.114 |
| iEEG | later | SI | -0.972 | 0.526 | 0.065 | 0.114 |
| iEEG | later | SI interactionbp | 605.250 | 130.059 | 0.000 | 0.000 |
| iEEG | later | SI secondslope | 604.278 | 129.916 | 0.000 | 0.000 |
| iEEG | later | DB | 0.786 | 0.477 | 0.099 | 0.114 |
| iEEG | later | PO | -0.616 | 0.495 | 0.213 | 0.213 |

Supplementary Table 9: Piecewise model outputs for Hippocampal Activity

| group | DV | IV | b | se | p | p FDR |
|---|---|---|---|---|---|---|
| Control | BOLD | SI | -0.008 | 0.004 | 0.024 | 0.024 |
| Control | BOLD | SI interactionbp | -0.169 | 0.045 | 0.000 | 0.000 |
| Control | BOLD | SI secondslope | -0.176 | 0.043 | 0.000 | 0.000 |
| Control | BOLD | DB | -0.016 | 0.004 | 0.000 | 0.000 |
| Control | BOLD | DB interactionbp | 0.035 | 0.007 | 0.000 | 0.000 |
| Control | BOLD | DB secondslope | 0.018 | 0.005 | 0.000 | 0.000 |
| Control | BOLD | PO | 0.015 | 0.003 | 0.000 | 0.000 |
| APOEe4 | BOLD | SI | -0.014 | 0.003 | 0.000 | 0.000 |
| APOEe4 | BOLD | DB | -0.017 | 0.005 | 0.000 | 0.000 |
| APOEe4 | BOLD | DB interactionbp | 0.028 | 0.007 | 0.000 | 0.000 |
| APOEe4 | BOLD | DB secondslope | 0.011 | 0.004 | 0.008 | 0.008 |
| APOEe4 | BOLD | PO | 0.014 | 0.003 | 0.000 | 0.000 |
| iEEG | theta | SI | 0.006 | 0.002 | 0.011 | 0.014 |
| iEEG | gamma | SI | -0.003 | 0.001 | 0.000 | 0.001 |
| iEEG | gamma | SI interactionbp | -0.854 | 0.261 | 0.001 | 0.004 |
| iEEG | gamma | SI secondslope | -0.857 | 0.261 | 0.001 | 0.001 |
| iEEG | theta | DB | -0.006 | 0.002 | 0.007 | 0.012 |
| iEEG | gamma | DB | 0.002 | 0.001 | 0.003 | 0.007 |
| iEEG | theta | PO | -0.005 | 0.002 | 0.018 | 0.018 |
| iEEG | gamma | PO | -0.005 | 0.002 | 0.015 | 0.017 |
| iEEG | gamma | PO interactionbp | 0.006 | 0.002 | 0.007 | 0.012 |
| iEEG | gamma | PO secondslope | 0.001 | 0.001 | 0.073 | 0.073 |

Supplementary Table 10: Piecewise model outputs for Hippocampal Activity - early vs late learning

| group | DV | learning phase | IV | b | se | p | p FDR |
|---|---|---|---|---|---|---|---|
| Control | BOLD | early | SI | -0.009 | 0.005 | 0.089 | 0.119 |
| Control | BOLD | early | DB | 0.000 | 0.005 | 0.928 | 0.928 |
| Control | BOLD | early | PO | 0.014 | 0.005 | 0.006 | 0.012 |
| Control | BOLD | early | Drop Error | 0.000 | 0.000 | 0.885 | 0.928 |
| Control | BOLD | late | SI | -0.007 | 0.004 | 0.089 | 0.119 |
| Control | BOLD | late | SI interactionbp | -0.237 | 0.052 | 0.000 | 0.000 |
| Control | BOLD | late | SI secondslope | -0.244 | 0.050 | 0.000 | 0.000 |
| Control | BOLD | late | DB | -0.019 | 0.004 | 0.000 | 0.000 |
| Control | BOLD | late | DB interactionbp | 0.039 | 0.008 | 0.000 | 0.000 |
| Control | BOLD | late | DB secondslope | 0.020 | 0.006 | 0.000 | 0.000 |
| Control | BOLD | late | PO | 0.016 | 0.003 | 0.000 | 0.000 |
| Control | BOLD | late | Drop Error | -0.001 | 0.001 | 0.013 | 0.022 |
| APOE | BOLD | early | SI | -0.017 | 0.006 | 0.004 | 0.010 |
| APOE | BOLD | early | DB | -0.027 | 0.011 | 0.009 | 0.013 |
| APOE | BOLD | early | DB interactionbp | 0.042 | 0.017 | 0.015 | 0.018 |
| APOE | BOLD | early | DB secondslope | 0.014 | 0.011 | 0.204 | 0.204 |
| APOE | BOLD | early | PO | 0.017 | 0.006 | 0.004 | 0.010 |
| APOE | BOLD | early | Drop Error | -0.001 | 0.000 | 0.154 | 0.154 |
| APOE | BOLD | late | SI | -0.013 | 0.003 | 0.000 | 0.001 |
| APOE | BOLD | late | DB | -0.014 | 0.005 | 0.008 | 0.013 |
| APOE | BOLD | late | DB interactionbp | 0.024 | 0.008 | 0.002 | 0.008 |
| APOE | BOLD | late | DB secondslope | 0.009 | 0.004 | 0.027 | 0.027 |
| APOE | BOLD | late | PO | 0.013 | 0.003 | 0.000 | 0.000 |
| APOE | BOLD | late | Drop Error | -0.001 | 0.000 | 0.009 | 0.013 |
| iEEG | HCtheta | early | SI | 0.003 | 0.003 | 0.226 | 0.289 |
| iEEG | HCgamma | early | SI | -0.004 | 0.001 | 0.000 | 0.000 |
| iEEG | HCtheta | early | DB | -0.004 | 0.003 | 0.122 | 0.178 |
| iEEG | HCgamma | early | DB | 0.002 | 0.001 | 0.006 | 0.021 |
| iEEG | HCtheta | early | PO | -0.004 | 0.003 | 0.130 | 0.178 |
| iEEG | HCgamma | early | PO | 0.001 | 0.001 | 0.312 | 0.358 |
| iEEG | HCtheta | early | Drop Error | 0.000 | 0.000 | 0.781 | 0.855 |
| iEEG | HCgamma | early | Drop Error | 0.000 | 0.000 | 0.007 | 0.021 |
| iEEG | HCtheta | late | SI | 0.014 | 0.005 | 0.003 | 0.021 |
| iEEG | HCgamma | late | SI | 0.000 | 0.001 | 0.997 | 0.997 |
| iEEG | HCgamma | late | SI interactionbp | -1.001 | 0.497 | 0.044 | 0.078 |
| iEEG | HCgamma | late | SI secondslope | -1.001 | 0.497 | 0.044 | 0.044 |
| iEEG | HCtheta | late | DB | -0.009 | 0.004 | 0.014 | 0.036 |
| iEEG | HCgamma | late | DB | -0.002 | 0.002 | 0.251 | 0.304 |
| iEEG | HCgamma | late | DB interactionbp | 0.010 | 0.004 | 0.013 | 0.036 |
| iEEG | HCgamma | late | DB secondslope | 0.008 | 0.003 | 0.009 | 0.009 |
| iEEG | HCtheta | late | PO | -0.008 | 0.004 | 0.040 | 0.076 |
| iEEG | HCgamma | late | PO | -0.012 | 0.004 | 0.006 | 0.021 |
| iEEG | HCgamma | late | PO interactionbp | 0.014 | 0.005 | 0.003 | 0.021 |
| iEEG | HCgamma | late | PO secondslope | 0.002 | 0.001 | 0.097 | 0.097 |
| iEEG | HCtheta | late | Drop Error | -0.001 | 0.000 | 0.086 | 0.142 |
| iEEG | HCgamma | late | Drop Error | 0.000 | 0.000 | 0.018 | 0.042 |

Supplementary Table 11: Piecewise model outputs for Hippocampal Activity- start vs end of trial

| group | DV | trial phase | IV | b | se | p | p FDR |
|---|---|---|---|---|---|---|---|
| Control | BOLD | start | SI | -0.001 | 0.005 | 0.918 | 0.918 |
| Control | BOLD | start | SI interactionbp | -0.180 | 0.065 | 0.005 | 0.009 |
| Control | BOLD | start | SI secondslope | -0.180 | 0.062 | 0.003 | 0.003 |
| Control | BOLD | end | SI | -0.012 | 0.005 | 0.008 | 0.012 |
| Control | BOLD | start | DB | -0.021 | 0.006 | 0.000 | 0.001 |
| Control | BOLD | start | DB interactionbp | 0.040 | 0.011 | 0.000 | 0.001 |
| Control | BOLD | start | DB secondslope | 0.019 | 0.008 | 0.021 | 0.021 |
| Control | BOLD | end | DB | -0.002 | 0.004 | 0.654 | 0.714 |
| Control | BOLD | start | PO | 0.015 | 0.004 | 0.000 | 0.001 |
| Control | BOLD | end | PO | 0.009 | 0.004 | 0.048 | 0.058 |
| Control | BOLD | start | Drop Error | -0.001 | 0.000 | 0.012 | 0.016 |
| Control | BOLD | end | Drop Error | 0.002 | 0.000 | 0.000 | 0.001 |
| APOE | BOLD | start | SI | -0.015 | 0.005 | 0.001 | 0.006 |
| APOE | BOLD | end | SI | -0.014 | 0.005 | 0.007 | 0.017 |
| APOE | BOLD | end | SI interactionbp | 0.505 | 0.235 | 0.032 | 0.053 |
| APOE | BOLD | end | SI secondslope | 0.491 | 0.233 | 0.035 | 0.035 |
| APOE | BOLD | start | DB | -0.016 | 0.007 | 0.033 | 0.053 |
| APOE | BOLD | start | DB interactionbp | 0.032 | 0.011 | 0.004 | 0.012 |
| APOE | BOLD | start | DB secondslope | 0.016 | 0.007 | 0.012 | 0.012 |
| APOE | BOLD | end | DB | -0.005 | 0.004 | 0.252 | 0.328 |
| APOE | BOLD | start | PO | -0.003 | 0.012 | 0.788 | 0.788 |
| APOE | BOLD | start | PO interactionbp | 0.033 | 0.016 | 0.042 | 0.060 |
| APOE | BOLD | start | PO secondslope | 0.030 | 0.007 | 0.000 | 0.000 |
| APOE | BOLD | end | PO | 0.012 | 0.005 | 0.013 | 0.028 |
| APOE | BOLD | start | Drop Error | -0.002 | 0.000 | 0.000 | 0.001 |
| APOE | BOLD | end | Drop Error | 0.000 | 0.000 | 0.326 | 0.385 |
| iEEG | HCtheta | start | SI | 0.001 | 0.005 | 0.821 | 0.864 |
| iEEG | HCgamma | start | SI | -0.005 | 0.002 | 0.011 | 0.227 |
| iEEG | HCtheta | end | SI | -0.005 | 0.005 | 0.287 | 0.575 |
| iEEG | HCgamma | end | SI | 0.001 | 0.002 | 0.537 | 0.716 |
| iEEG | HCtheta | start | DB | -0.010 | 0.005 | 0.059 | 0.328 |
| iEEG | HCgamma | start | DB | 0.002 | 0.002 | 0.441 | 0.678 |
| iEEG | HCtheta | end | DB | -0.008 | 0.005 | 0.120 | 0.328 |
| iEEG | HCgamma | end | DB | -0.001 | 0.002 | 0.602 | 0.752 |
| iEEG | HCtheta | start | PO | -0.007 | 0.005 | 0.147 | 0.328 |
| iEEG | HCgamma | start | PO | 0.003 | 0.002 | 0.105 | 0.328 |
| iEEG | HCtheta | end | PO | -0.003 | 0.005 | 0.528 | 0.716 |
| iEEG | HCgamma | end | PO | -0.001 | 0.002 | 0.771 | 0.864 |
| iEEG | HCtheta | start | Drop Error | 0.000 | 0.000 | 0.987 | 0.987 |
| iEEG | HCgamma | start | Drop Error | 0.000 | 0.000 | 0.098 | 0.328 |
| iEEG | HCtheta | end | Drop Error | 0.000 | 0.000 | 0.378 | 0.676 |
| iEEG | HCgamma | end | Drop Error | 0.000 | 0.000 | 0.139 | 0.328 |

Supplementary Table 12: Theta Power Effects by Frequency Band

| frequency | IV | p |
|---|---|---|
| 3Hz | SI | 0.006 |
| 4Hz | SI | 0.015 |
| 5Hz | SI | 0.022 |
| 6Hz | SI | 0.093 |
| 7Hz | SI | 0.267 |
| 8Hz | SI | 0.06 |
| 3Hz | DB | 0.022 |
| 4Hz | DB | 0.049 |
| 5Hz | DB | 0.076 |
| 6Hz | DB | 0.09 |
| 7Hz | DB | 0.016 |
| 8Hz | DB | 0.004 |
| 3Hz | PO | 0.01 |
| 4Hz | PO | 0.082 |
| 5Hz | PO | 0.061 |
| 6Hz | PO | 0.105 |
| 7Hz | PO | 0.22 |
| 8Hz | PO | 0.052 |
| 3Hz | Drop Error | 0.975 |
| 4Hz | Drop Error | 0.636 |
| 5Hz | Drop Error | 0.254 |
| 6Hz | Drop Error | 0.272 |
| 7Hz | Drop Error | 0.797 |
| 8Hz | Drop Error | 0.267 |

Supplementary Table 13: Piecewise model outputs for Entorhinal and Caudate Activity

| group | region | IV | b | se | p | p FDR |
|---|---|---|---|---|---|---|
| Control | Entorhinal | SI | -0.019 | 0.004 | 0.000 | 0.000 |
| Control | Entorhinal | DB | -0.014 | 0.006 | 0.014 | 0.026 |
| Control | Entorhinal | DB interactionbp | 0.043 | 0.010 | 0.000 | 0.000 |
| Control | Entorhinal | DB secondslope | 0.029 | 0.007 | 0.000 | 0.000 |
| Control | Entorhinal | PO | 0.018 | 0.004 | 0.000 | 0.000 |
| Control | Entorhinal | Drop Error | 0.000 | 0.000 | 0.372 | 0.465 |
| Control | Caudate | SI | 0.008 | 0.005 | 0.141 | 0.201 |
| Control | Caudate | SI interactionbp | -0.252 | 0.070 | 0.000 | 0.001 |
| Control | Caudate | SI secondslope | -0.244 | 0.067 | 0.000 | 0.000 |
| Control | Caudate | DB | 0.010 | 0.004 | 0.016 | 0.026 |
| Control | Caudate | PO | 0.001 | 0.004 | 0.827 | 0.880 |
| Control | Caudate | Drop Error | 0.000 | 0.000 | 0.880 | 0.880 |
| APOEe4 | Entorhinal | SI | -0.021 | 0.004 | 0.000 | 0.000 |
| APOEe4 | Entorhinal | DB | -0.023 | 0.007 | 0.001 | 0.003 |
| APOEe4 | Entorhinal | DB interactionbp | 0.040 | 0.010 | 0.000 | 0.000 |
| APOEe4 | Entorhinal | DB secondslope | 0.017 | 0.006 | 0.004 | 0.004 |
| APOEe4 | Entorhinal | PO | 0.016 | 0.004 | 0.000 | 0.000 |
| APOEe4 | Entorhinal | Drop Error | 0.000 | 0.000 | 0.476 | 0.612 |
| APOEe4 | Caudate | SI | -0.002 | 0.004 | 0.604 | 0.680 |
| APOEe4 | Caudate | DB | -0.005 | 0.004 | 0.233 | 0.350 |
| APOEe4 | Caudate | PO | 0.000 | 0.004 | 0.923 | 0.923 |
| APOEe4 | Caudate | Drop Error | -0.001 | 0.000 | 0.016 | 0.030 |
